# PKC isoforms activate LRRK1 kinase by phosphorylating conserved residues (Ser1064, Ser1074 and Thr1075) within the COR_B_ GTPase domain

**DOI:** 10.1101/2022.06.09.495448

**Authors:** Asad U Malik, Athanasios Karapetsas, Raja S. Nirujogi, Deep Chatterjee, Toan K. Phung, Melanie Wightman, Robert Gourlay, Nick Morrice, Sebastian Mathea, Stefan Knapp, Dario R Alessi

## Abstract

Leucine-rich-repeat-kinase 1 (LRRK1) and its homologue LRRK2 are multidomain kinases possessing a ROC-COR_A_-COR_B_ containing GTPase domain and phosphorylate distinct Rab proteins. LRRK1 loss of function mutations cause the bone disorder osteosclerotic metaphyseal dysplasia, whereas LRRK2 missense mutations that enhance kinase activity cause Parkinson’s disease. Previous work suggested that LRRK1 but not LRRK2, is activated via a Protein Kinase C (PKC)-dependent mechanism. Here we demonstrate that phosphorylation and activation of LRRK1 in HEK293 cells is blocked by PKC inhibitors including LXS-196 (Darovasertib), a compound that has entered clinical trials. We show multiple PKC isoforms phosphorylate and activate recombinant LRRK1 in a manner reversed by phosphatase treatment. PKCα unexpectedly does not activate LRRK1 by phosphorylating the kinase domain, but instead phosphorylates a cluster of conserved residues (Ser1064, Ser1074 and Thr1075) located within a region of the COR_B_ domain of the GTPase domain. These residues are positioned at the equivalent region of the LRRK2 DK helix reported to stabilize the kinase domain αC-helix in the active conformation. Thr1075 represents an optimal PKC site phosphorylation motif and its mutation to Ala, blocked PKC-mediated activation of LRRK1. A triple Glu mutation of Ser1064/Ser1074/Thr1075 to mimic phosphorylation, enhanced LRRK1 kinase activity ~3-fold. From analysis of available structures, we postulate that phosphorylation of Ser1064, Ser1074 and Thr1075 activates LRRK1 by promoting interaction and stabilization of the aC-helix on the kinase domain. This study provides new fundamental insights into the mechanism controlling LRRK1 activity and reveals a novel unexpected activation mechanism.

## Introduction

The Leucine Rich Repeat Protein Kinase-1 (LRRK1) and its paralog LRRK2 are large multidomain protein kinases possessing ANK, LRR, ROC-COR_A_-COR_B_ containing ROCO GTPase, kinase and WD40 domains [1]. In addition, LRRK2 possesses an N-terminal ARM domain not-present in LRRK1 [2]. Rare recessive loss of function mutations in LRRK1 lead to a serious bone disorder termed osteosclerotic metaphyseal dysplasia [3, 4]. Autosomal dominant mutations that enhance the kinase activity of LRRK2 cause familial late onset Parkinson’s that is similar in phenotype to the idiopathic form of the disease [5, 6]. LRRK1 and LRRK2 phosphorylate distinct Rab GTPase proteins within their Switch-II effector binding motif. LRRK1 phosphorylates Rab7A at Ser72 [7, 8], whereas LRRK2 phosphorylates a subset of Rab proteins, including Rab8A, Rab10 and Rab12 [9, 10]. LRRK2 phosphorylation of Rab proteins does not affect GTPase activity but controls effector binding by promoting interaction with effectors such as RILPL1/RILPL2 [10,Dhekne, 2018 #5728, 11], JIP3/JIP4 [11, 12] and myosin-Va [13] that are critical in controlling downstream biology.

Our previous work suggests that LRRK1, but not LRRK2, is stimulated by phorbol esters that switch on conventional and novel PKC isoforms [8]. In contrast, LRRK2 is activated by recruitment to membrane vesicles, through binding to Rab GTPases such as Rab29 that interacts with the ARM domain that is not present in LRRK1 [14–18]. Furthermore, a Parkinson’s causing mutation in VPS35[D620N] stimulates LRRK2 activity without impacting LRRK1 [8, 19]. Thus, although LRRK1 and LRRK2 are close homologues, mutations in these two kinases have been linked to the development of diverse diseases, they are stimulated by distinct pathways and phosphorylate different Rab substrates (Table 1).

**Table 1:** Comparing LRRK1 and LRRK2.

|  |  |  |
| --- | --- | --- |
|  | **LRRK1** | **LRRK2** |
| **Associated disease** | Osteosclerotic metaphyseal dysplasia (loss of function mutation) [3, 4] | Parkinson's disease (gain of function/kinase domain activating mutations) [5, 6]. |
| **Substrates** | Rab7A [7, 8]. LRRK1 does not phosphorylate Rab8A and Rab10, further work needed to investigate other Rab targets | Rab1, Rab3, Rab8, Rab10, Rab12, Rab29, Rab35 and Rab43. [9, 10]. LRRK2 does not phosphorylate Rab7A [8] |
| **Upstream regulation** | Phorbol ester or EGF stimulation [8] | 1. Recruitment of LRRK2 to membranes by binding to Rab proteins (e.g Rab8A, Rab10 and Rab29 and pRab8A, pRab10) via ARM domain [14, 15, 17].<br>2. Parkinson's mutations of the retromer complex [19].<br>3. Lysosomal stress/damage candida infection, LLOMe, chloroquine [12, 55, 56]<br>4. Nigericin and monensin (unknown mechanism) [57] |

There are multiple isoforms of PKC belonging to the larger family of AGC protein kinases [20, 21]. Many AGC kinases including all PKC isoforms are activated by T-loop phosphorylation by the 3-phosphoinositide dependent protein kinase-1 (PDK1) [22, 23]. Activity of conventional and novel PKC isoforms is also regulated by phosphorylation of their hydrophobic motif residue that recent work suggests is controlled by autophosphorylation via the mTORC2 kinase phosphorylating the TOR Interacting motif [24]. Conventional (PKCα, PKCβ, PKCγ) and novel (PKCδ, PKCε, PKCη and PKCθ) PKC isoforms are stimulated by lipids such as diacylglycerol though only conventional isoforms also bind and are activated by Ca^2+^ [25]. Phorbol esters are substances from plants that mimic the function of diacylglycerol. They are widely used tools to activate PKCs in cell-culture experiments. The atypical PKCs (PKC*i* and PKCz) are regulated by a distinct upstream pathway but still require PDK1 to be activated [26–28]. PKC isoforms phosphorylate a multitude of substrates involved in regulating a wide range of cellular pathways. Classical PKC substrates are phosphorylated on a Ser/Thr residue lying in a motif that possess a basic residue at the –3 and –5 positions and a hydrophobic residue at the + 1 positions [29], although PKCs can also phosphorylate substrates at non canonical sites deviating from this optimal motif [30].

We previously observed that stimulation of mouse embryonic fibroblasts (MEFs) with phorbol ester, induced marked phosphorylation of Rab7A in a manner that was blocked by knock-out of LRRK1, treatment with a non-specific LRRK1 inhibitor GZD824 or treatment with a relatively non-specific GÖ6983 PKC isoform inhibitor [8]. These results suggested that PKC isoforms play a role in regulating the activation of LRRK1, but the mechanism by which PKC could control LRRK1 remained elusive. In this study we explored the activation mechanism further and demonstrate that PKC isoforms directly activate LRRK1 by phosphorylating a cluster of highly conserved residues (Ser1064, Ser1074 and Ser1075) located within the COR_B_ domain that forms part of the Roco GTPase domain. Our data provides firm evidence that LRRK1 is a direct downstream target of PKC isoforms. After PKD isoforms that are also activated by PKC isoforms [31], LRRK1 is thus the second class of kinase to be demonstrated to be directly activated by PKC isoforms. It also establishes a new paradigm by which phosphorylation of the COR_B_ located within the GTPase domain can directly stimulate LRRK1 kinase activity.

## Results

### Establishing a cell system to dissect the mechanism of LRRK1 regulation by PKC isoforms

Consistent with previous findings in mouse embryonic fibroblasts [32], phorbol ester stimulation enhanced Rab7A phosphorylation at Ser72 in control non-transfected HEK293 cells, and to greater extent in HEK293 cells that stably overexpress GFP-LRRK1 (Fig 1A). This was blocked by the PKC inhibitor GÖ6983 (1 μM) (Fig 1A). Immunoblotting with a pan selective PKC site phospho-specific substrate antibody [33], confirmed phorbol ester triggered phosphorylation of a multitude of cellular proteins which was suppressed by GÖ6983 (Fig 1A). A time course of phorbol ester stimulation revealed that increased Rab7A phosphorylation was detectable within 1 min, plateauing between 20 and 40 min, and declining thereafter, paralleling phosphorylation of ERK1/2 (Fig 1B, quantified in SFig 1), that is triggered by PKC activation [34]. PKCα levels decreased ~40% between 80 to 160 min paralleling a similar decline in Rab7A phosphorylation (Fig 1B, quantified in SFig 1). Stable expression of kinase inactive LRRK1[D1409A], did not increase Rab7A phosphorylation beyond that induced by endogenous LRRK1 following phorbol ester treatment (Fig 1C). Consistent with LRRK2 not being controlled by PKC, phosphorylation of the LRRK2 specific substrate Rab10 at Thr73, was not impacted by phorbol ester stimulation (Fig 1C). Bryostatin-1 a macrocyclic lactone produced by the marine organism *Bugula neritina* that binds to the phorbol ester receptor and activates PKC isoforms [35], also triggered phosphorylation of Rab7A in a manner that was blocked by GÖ6983 (Fig 1D).

**Figure 1.**
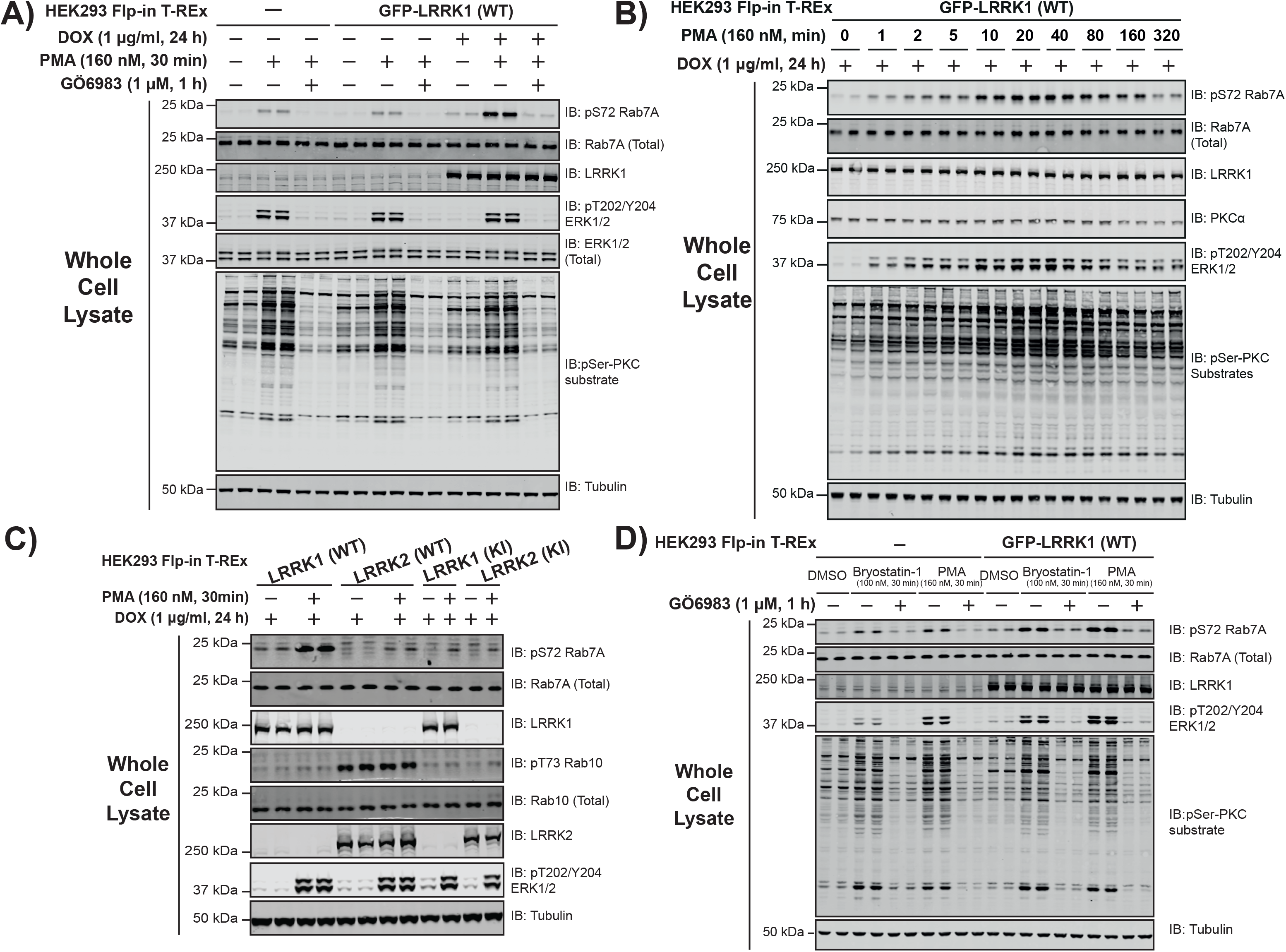
Phorbol ester induces Rab7A phosphorylation in HEK293 Flp-In T-REx cells. **(A)** HEK293 Flp-In T-REx cells stably expressing wild type (WT) GFP-LRRK1, were treated ± 1 μg/ml doxycycline for 24 h to induce expression of GFP-LRRK1. Cells were serum-starved for 16 h, incubated ± 1 μM GÖ6983 for 1 h and 30 min prior to cell lysis were stimulated ± 160 nM phorbol 12-myristate 13-acetate (PMA) for 30 min. 10 μg of cell extracts were subjected to quantitative immunoblot analysis using the LI-COR Odyssey CLx Western Blot imaging system with the indicated antibodies. Combined immunoblotting data from 2 independent biological replicates (each performed in duplicate) are shown. **(B)** Top panel-as in **(A)** except cells were stimulated with PMA for the indicated timepoints. Lower panel quantified immunoblotting data are presented as ratios of pRab7ASer72/total Rab7A (mean ± SEM). **(C)** As in **(A)** except cells that induce express GFP-LRRK1[D1409A] (kinase-inactive), GFP-LRRK2 WT or GFP-LRRK2[D2017A] (kinase-inactive) are also employed. **(D)** As in **(A)** except cells were incubated ± 1 μM GÖ6983 for 1 h and 30 min prior to cell lysis were stimulated with either ± 100 nM Bryostatin-1 or 160 nM PMA for 30 min.

### Optimizing cell lysis conditions to preserve LRRK1 kinase activity

In performing pilot immunoprecipitation studies, we noted that immunoprecipitated LRRK1 isolated from cell lysates prepared in 1% (v/v) Triton-X100 detergent, displayed low activity when compared with recombinant LRRK1[20-2015] produced in insect cells (Fig 2A). This prompted us to investigate other detergents and we found that the kinase activity of immunoprecipitated LRRK1 activity was significantly higher and closer to the activity of recombinant LRRK1[20-2015] when cells were lysed with subcritical concentrations of weak non-ionic detergents 1% (w/v) digitonin or Zwitterionic 0.3% (w/v) CHAPS (Fig 2A). In all subsequent studies in which LRRK1 is immunoprecipitated, cells were lysed with 0.3% (w/v) CHAPS (mainly due to its lower cost compared to digitonin). We also noted that levels of LRRK1 that were recovered were similar with all detergents, however, ~50% lower levels of pRab7A and total Rab7A, were solubilized in cell extracts prepared with 1% (w/v) digitonin or 0.3% (w/v) CHAPS compared with 1% (v/v) Triton-X100 (Fig 2B). We also investigated whether supplementing the cell lysis buffer with 10 μM GTP-γ-S and 1 mM MgCl_2_ to preserve GTP binding to the ROC GTPase domain impacted kinase activity but observed that this treatment did not significantly alter the kinase activity in *in vitro* kinase assays (Fig 2A).

**Figure 2.**
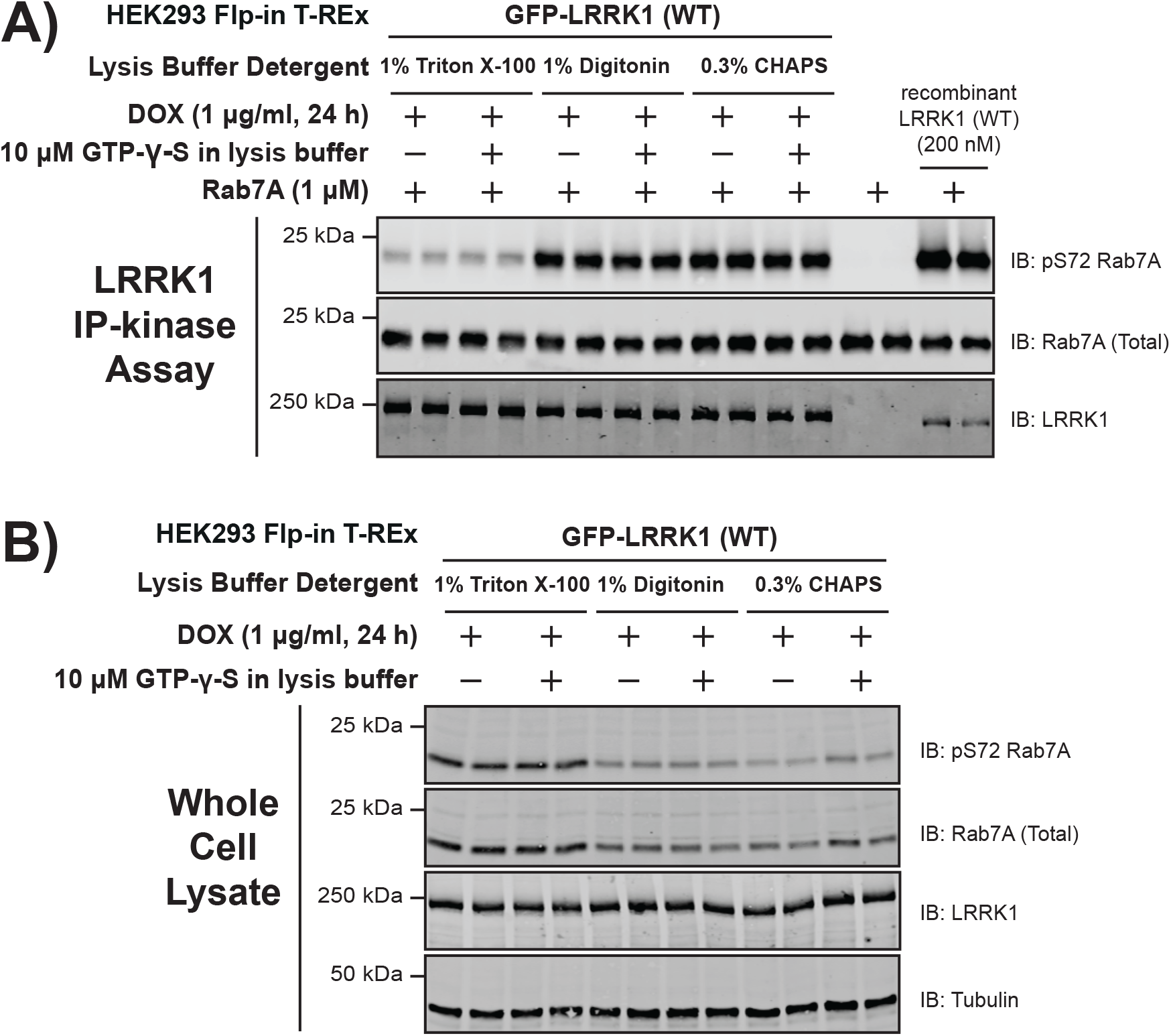
Optimizing cell lysis conditions to preserve LRRK1 kinase activity. (**A**) HEK293 Flp-In T-REx cells stably expressing wild type (WT) GFP-LRRK1, were treated with 1 μg/ml doxycycline for 24 h to induce expression of GFP-LRRK1. Cells were harvested in a lysis buffer containing (50 mM HEPES, pH 7.4, 270 mM sucrose, 150 mM NaCl, 1 mM sodium orthovanadate, 50 mM NaF, 10 mM 2-glycerophosphate, 5 mM sodium pyrophosphate, 1 μg/ml microcystin-LR and complete EDTA-free protease inhibitor cocktail) the indicated detergent ± 10 μM GTP-γ-S. GFP-LRRK1 was immunoprecipitated from cell lysates using an anti-GFP nanobody. Immunoprecipitates were washed in the same cell lysis buffer before being washed into kinase assay buffer (25 mM HEPES pH7.4, 50 mM KCl, 0.1% (v/v 2-mercaptoethanol). LRRK1 kinase activity towards recombinant Rab7A was assessed in a 30 min assay. Reactions were terminated by addition of SDS-sample buffer and levels of pSer72-Rab7A, Rab7A and LRRK1 were assessed by quantitative immunoblot analysis using the LI-COR Odyssey CLx Western Blot imaging system with the indicated antibodies. Combined immunoblotting data from 2 independent biological replicates (each performed in duplicate) are shown. (**B**) The whole cell extracts (10 μg) that were prepared in (**A**) were subjected to quantitative immunoblot analysis using the LI-COR Odyssey CLx Western Blot imaging system with the indicated antibodies.

### PKC isoforms phosphorylate and activate LRRK1-in cells

We next stimulated cells ± phorbol ester for 30 min, immunoprecipitated LRRK1 and assayed its activity *in vitro* employing Rab7A as a substrate. This revealed that phorbol ester stimulation markedly enhanced activity of immunoprecipitated LRRK1 by ~7.5-fold (Fig 3A). Immunoblotting revealed that phorbol ester increased recognition of immunoprecipitated LRRK1 with a pan-phospho-PKC substrate motif antibody [36], consistent with the notion that PKC directly phosphorylates LRRK1 (Fig 3A). Incubation of cells with increasing doses of the LXS-196 (Darovasertib), a PKC inhibitor, that is currently in human clinical cancer trials [37], and highly specific [38], resulted in a concentration dependent inhibition of immunoprecipitated LRRK1 kinase activity, with an IC50 of ~40 nM (Fig 3A). The inhibition of LRRK1 activity in vitro paralleled that of endogenous Rab7A phosphorylation, as well as cellular PKC substrate phosphorylation (Fig 3B). LXS-196 is structurally unrelated to GÖ6983 that is a staurosporine analogue and a much less selective kinase inhibitor [39]. The GÖ6983 inhibitor also blocked phorbol ester induced activation of LRRK1 activity (Fig 3B).

**Figure 3.**
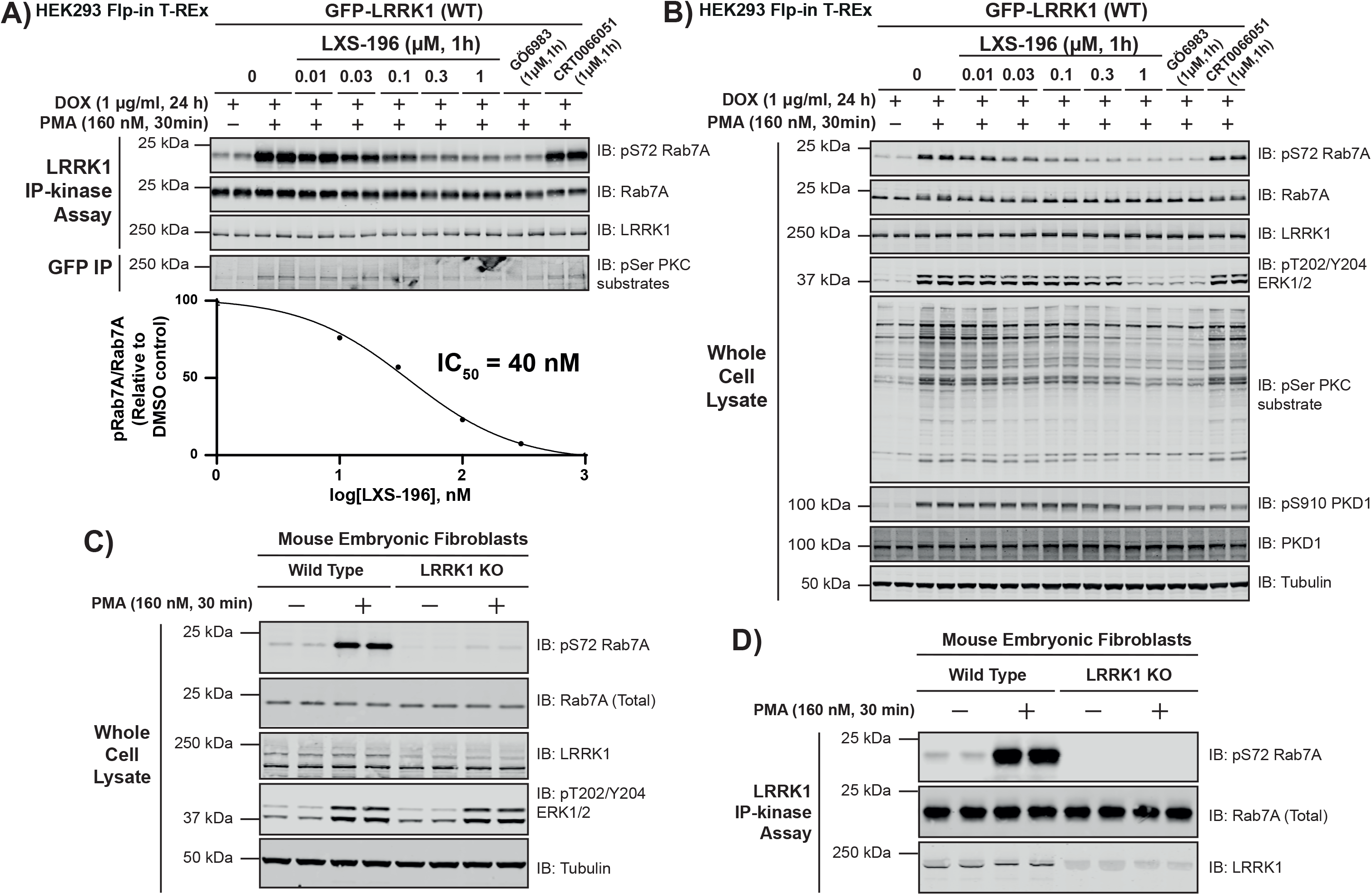
PMA induces phosphorylation and activation of immunoprecipitated LRRK1. **(A, upper panel)** HEK293 Flp-In T-REx cells stably expressing wild type (WT) GFP-LRRK1 were treated with 1 μg/ml doxycycline for 24 h to induce expression of GFP-LRRK1. Cells were then serum-starved for 16 h, incubated ± the indicated concentration of LXS-196, GÖ6983 or CRT0066051. 30 min prior to lysis, cells were stimulated ± 160 nM phorbol 12-myristate 13-acetate (PMA) for 30 min and cell lysed. Upper panel-GFP-LRRK1 immunoprecipitated using an anti-GFP nanobody and LRRK1 kinase activity towards recombinant Rab7A assessed in a 30 min assay. Reactions were terminated by addition of SDS-sample buffer and levels of pSer72-Rab7A, Rab7A and LRRK1 were assessed by quantitative immunoblot analysis using the LI-COR Odyssey CLx Western Blot imaging system with the indicated antibodies. Combined immunoblotting data from 2 independent biological replicates (each performed in duplicate) are shown. **(A, lower panel)** Quantified immunoblotting data of LRRK1 kinase assays are presented as ratios of pRab7ASer72/total Rab7A (mean ± SEM) vs concentration of LXS-196 relative to levels observed with no inhibitor added (100%). IC50 values were calculated with GraphPad Prism (version 9.1.0) using non-linear regression analysis. Note that the blank value of basal LRRK1 activity observed in unstimulated cells with no inhibitor was subtracted from stimulated values for the IC50 analysis (**B**) The whole cell extracts (10 μg) that were prepared in (**A**) were subjected to quantitative immunoblot analysis using the LI-COR Odyssey CLx Western Blot imaging system with the indicated antibodies. **(C)** As in **(A)** except wild type and homozygous LRRK1 knock-out primary Mouse Embryonic Fibroblasts (MEFs) were serum-starved for 16 h and stimulated ± 160 nM PMA for 30 min and cells lysed. Whole cell extracts (10 μg) were prepared and subject to quantitative immunoblot analysis. **(D)** As in **(C)** except, endogenous LRRK1 was immunoprecipitated from whole cell extracts using a polyclonal total LRRK1 antibody and LRRK1 kinase activity towards recombinant Rab7A assessed in a 30 min assay. Phosphorylation off Rab7A was assessed by immunoblot analysis.

PKC isoforms activate PKD kinases [31, 40]. To rule out PKD activating LRRK1, we used a PKD kinase inhibitor termed CRT0066051 (1 μM) [41] and found that treating cells with this selective PKD inhibitor did not impact phorbol ester mediated activation of LRRK1 (Fig 3B). CRT0066051 reduced autophosphorylation of PKD1 at Ser910 confirming that inhibitor suppressed kinase PKD1 activity (Fig 3B) [41]. As expected, LXS-196 and GÖ6983 PKC inhibitors also suppressed autophosphorylation of PKD1 at Ser910 (Fig 3B).

We investigated whether the kinase activity of endogenous LRRK1 was similarly stimulated upon phorbol ester treatment. To this end, we employed previously characterized wild type and homozygous LRRK1 knock-out mouse embryonic fibroblasts [8] and stimulated these cells ± phorbol ester. Endogenous LRRK1 was immunoprecipitated and assayed employing recombinant Rab7A as a substrate. This revealed that phorbol ester markedly stimulated endogenous LRRK1 kinase activity and Rab7A phosphorylation in wild type but not in LRRK1 knockout cells (Fig 3C, 3D).

### PKC isoforms phosphorylate and activate LRRK1-*in vitro*

We next tested whether recombinant PKC isoforms (PKCα, PKCβ, PKCγ, PKCε, PKCθ and PKCζ) directly phosphorylated wild type LRRK1[20-2015] and kinase inactive LRRK1[20-2015, D1409A]. Reactions were undertaken using a Mg-[γ-^32^P]ATP radioactive assay and phosphorylation monitored by autoradiography as well as by immunoblotting with the Pan phospho-PKC substrate antibody (Fig 4A). All PKC isoforms induced phosphorylation of wild type as well as kinase-inactive LRRK1[D1409A] to differing extents, with PKCα, PKCβ, and PKCθ phosphorylating LRRK1 most strongly. Autophosphorylation of all PKC isoforms except for PKCγ was observed (Fig 4A). Although PKCγ was not detected in the reaction mixture it was observed in a Coomassie stained gel of the PKC isoforms panel used for this study (Fig 4B). It is possible that PKCγ became degraded during the kinase assay in this experiment. Additionally, in reactions employing PKCζ, a protein band that is likely a phosphorylated contaminant in the PKCζ preparation, is visible below the ~250 kDa LRRK1 band on the autoradiograph. Using Mg-[γ-^32^P]ATP of known specific activity, we were able to estimate the stoichiometry of PKCα mediated phosphorylation of LRRK1. At the highest concentration of PKCα tested (100 nM), stoichiometry of phosphorylation was calculated to be 0.23 mol ^32^P/mol protein (SFig 2).

**Figure 4.**
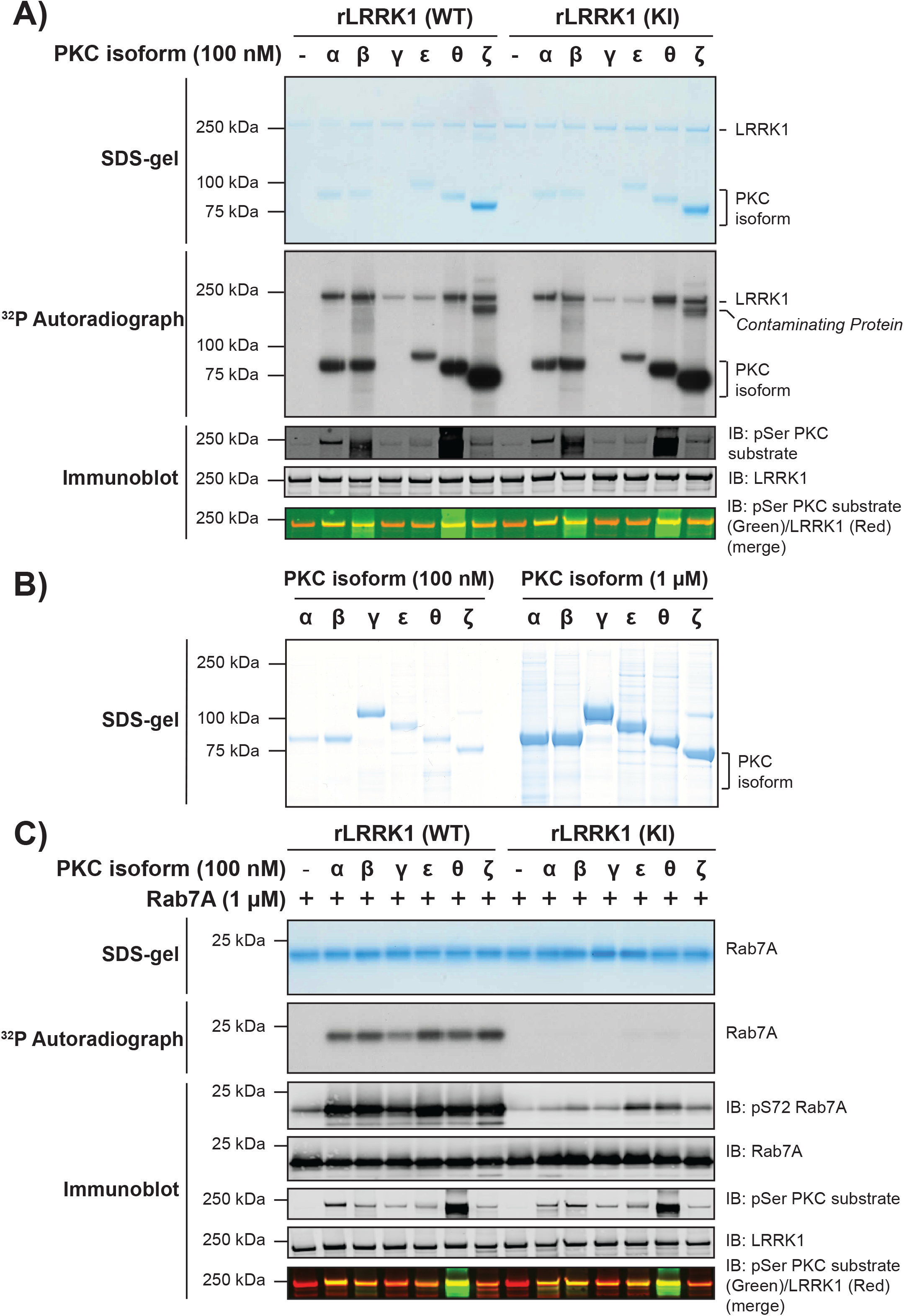
Recombinant PKC isoforms phosphorylate and activate LRRK1. **(A)** The indicated recombinant PKC isoforms (100 nM) were incubated with recombinant wild type (WT) insect cell expressed recombinant (r) LRRK1[20-2015] or kinase inactive (KI) LRRK1[D1409A, 20-2015] (50 nM) in the presence of Mg[γ-^32^P]ATP. Reactions were terminated after 30 min with SDS-sample buffer. 80% of each reaction was resolved by SDS-polyacrylamide electrophoresis, stained by Coomassie blue (upper panel), and subjected to autoradiography (middle panel). The remaining reaction mixture was subjected to a multiplexed immunoblot analysis using the LI-COR Odyssey CLx Western Blot imaging system to assess pan PKC substrate phosphorylation antibody (green channel) and total levels of LRRK1 protein (red channel) (lower panel). **(B)** 20 μl of 100 nM (left panel) or 1 μM (right panel), of the indicated PKC isoforms was used in panel (A) were subjected to polyacrylamide gel electrophoresis and stained with Coomassie blue. **(C)** The indicated recombinant PKC isoforms (100 nM) were incubated with recombinant wild type LRRK1[20-2015] or kinase inactive LRRK1[D1409A, 20-2015] (50 nM) in the presence of non-radioactive MgATP for 30 min (Step-1 kinase assay). PKC phosphorylated LRRK1 was then diluted 1.5-fold into a kinase assay containing recombinant Rab7A (1 mM) in the presence of Mg[γ-^32^P]ATP (Step 2 of the kinase assay). Reactions were terminated after 30 min with SDS-sample buffer. 75% of each reaction was subjected to SDS-polyacrylamide electrophoresis, stained by Coomassie blue (upper panel), and subjected to autoradiography (middle panel). The remaining reaction mixture was subjected to a multiplexed immunoblot analysis using the LI-COR Odyssey CLx Western Blot imaging system with the indicated antibodies (lower panel).

To determine whether PKC phosphorylation of LRRK1 stimulated kinase activity, we performed a two-step kinase assay. In the first step, recombinant wild type LRRK1[20-2015] or kinase-inactive LRRK1[20-2015, D1409A], was incubated for 30 min with PKC isoforms (PKCα, PKCβ, PKCγ, PKCε, PKCθ and PKCζ) in the presence of non-radioactive Mg-ATP. In the second step, an aliquot of this first reaction was assayed for its ability to phosphorylate Rab7A in the presence of Mg[γ-^32^P]ATP. Phosphorylation of Rab7A was monitored by ^32^P-autoradiography as well as with a pSer72 Rab7A phospho-specific antibody. We observed that PKC phosphorylation of wild type LRRK1[20-2015], but not kinase-inactive LRRK1[20-2015, D1409A], markedly boosted phosphorylation of Rab7A, with all PKC isoforms tested (Fig 4C). It should be noted that low levels of Rab7A phosphorylation at Ser72 was detected with PKCε and PKCθ phosphorylation of kinase-inactive LRRK1[20-2015, D1409A] (Fig 4C), which is likely due to these PKC isoforms phosphorylating Rab7A to a very low stoichiometry, which can be detected using the high sensitivity phospho-specific antibody.

We next analyzed whether wild type LRRK1 immunoprecipitated from unstimulated HEK293 cells could be activated by PKCα phosphorylation in vitro. For these studies we used PKCα as this is one of the most well studied isoforms and we were able to obtain sufficient enzyme and a high concentration for our analysis. We treated immunoprecipitated LRRK1 with increasing doses of recombinant PKCα (10 to 300 nM) for 30 min in the presence of Mg^2+^ ATP and subsequently measured phosphorylation of Rab7A employing a phospho-specific antibody (Fig 5A). Increased Rab7A phosphorylation was detected in this assay using 30 nM PKCα, plateauing at an enzyme concentration of 100 nM. Similar results were obtained from an equivalent experiment in which recombinant LRRK1[20-2015] produced in insect cells was phosphorylated with increasing amounts of PKCα (SFig 3A). We next preformed a time course of activation of LRRK1 using 100 nM PKCα and found that LRRK1 activation was detected after 5 min incubation and plateaued at 40 min (Fig 5B). Similar results were obtained in an analogous experiment employing insect cell-produced recombinant LRRK1[20-2015] (SFig 3B). We also found that the activity of phorbol ester-stimulated LRRK1 was moderately enhanced by PKCα phosphorylation in vitro (SFig 4).

**Figure 5.**
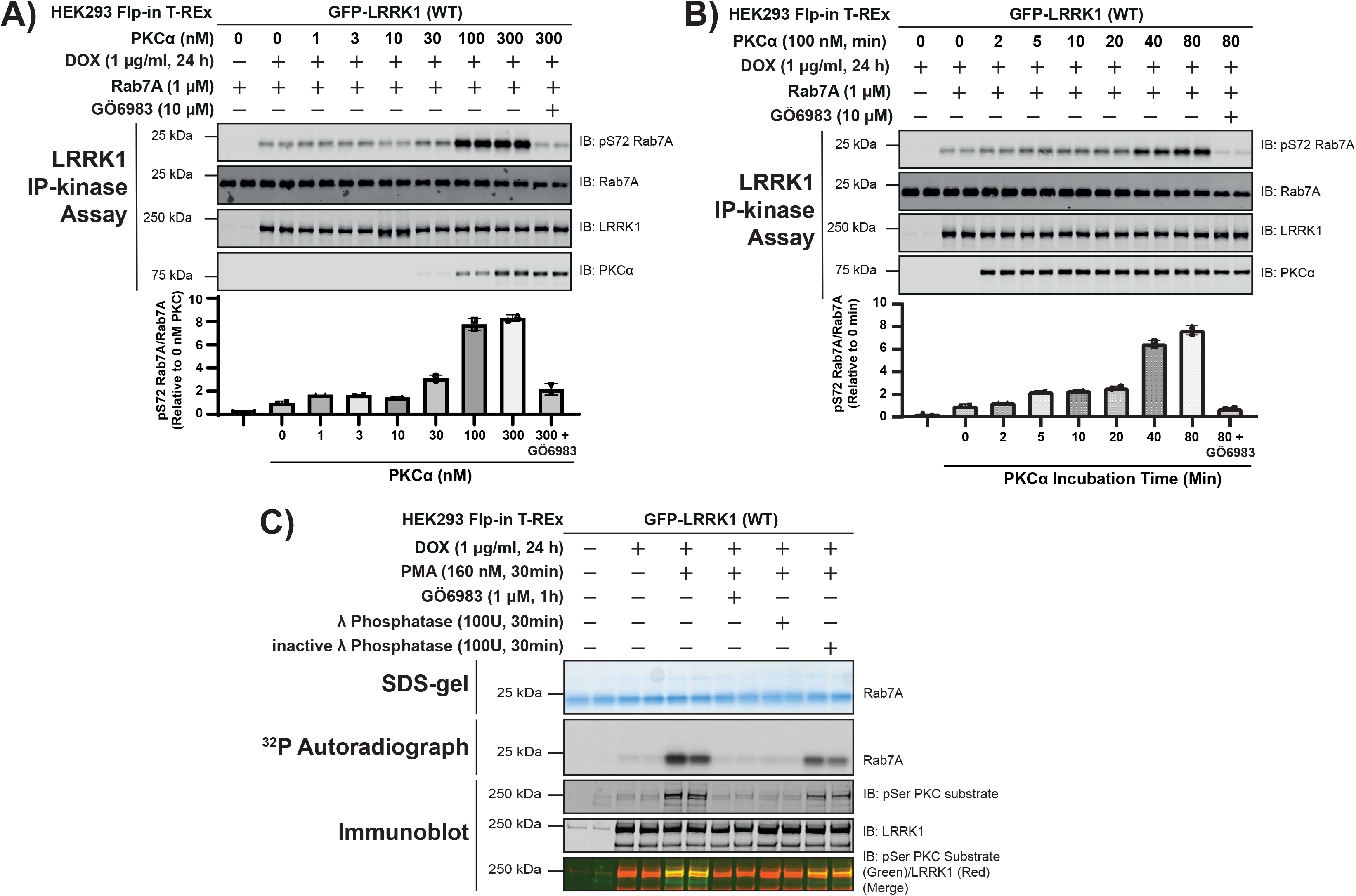
Time-course, dose-dependence, and reversibility of activation of LRRK1 by PKCα. **(A)** HEK293 Flp-In T-REx cells stably expressing wild type (WT) GFP-LRRK1 were treated ± 1 μg/ml doxycycline for 24 h to induce GFP-LRRK1 expression. Cells were serum-starved for 16 h, lysed and GFP-LRRK1 immunoprecipitated and aliquoted as indicated. Each aliquot contains GFP-LRRK1 immunoprecipitated from 1 mg of HEK293 cell lysate. Step-1 of the LRRK1 kinase activation assay was setup by incubating GFP-LRRK1 aliquots with the indicated concentrations of PKCα and ± inhibitor GÖ6983 (10 μM) in the presence of non-radioactive MgATP for 30 min. PKC phosphorylated LRRK1 was then diluted 1.5-fold into a kinase assay containing recombinant Rab7A (1 mM) in the presence of non-radioactive MgATP (Step 2 of the kinase assay). Reactions were terminated after 30 min with SDS-sample buffer analysed by multiplexed immunoblot analysis using the LI-COR Odyssey CLx Western Blot imaging system with the indicated antibodies (upper panel). Combined immunoblotting data from 2 independent biological replicates (each performed in duplicate) are shown. Lower panel quantified immunoblotting data are presented as ratios of pRab7ASer72/total Rab7A (mean ± SEM) relative to levels observed with no PKCα added (given a value of 1.0). (**B**) As in (**A**) except in step 1 of the kinase activity assay 100 nM PKCα was incubated with immunoprecipitated GFP-LRRK1 for the times indicated. (**C**) As in (A) except that following serum starvation, cells were treated stimulated ± 160 nM phorbol 12-myristate 13-acetate (PMA) for 30 min and cells lysed. GFP-LRRK1 was immunoprecipitated from 1 mg of cell lysate incubated ± 100U of either active or 10 mM EDTA-inactivated λ phosphatase for 30 min. Following the phosphatase treatment, the beads were extensively washed and LRRK1 was subjected to kinase assay with recombinant Rab7A using Mg[γ-^32^P]ATP. 75% of each reaction was subjected to SDS-polyacrylamide electrophoresis, stained by Coomassie blue (upper panel), and subjected to autoradiography (middle panel). The remaining reaction mixture was subjected to a multiplexed immunoblot analysis using the LI-COR Odyssey CLx Western Blot imaging system with the indicated antibodies (lower panels). Combined immunoblotting data from 2 independent biological replicates (each performed in duplicate) are shown.

Consistent with phosphorylation triggering activation of LRRK1, incubation of immunoprecipitated activated LRRK1 derived from phorbol ester stimulated cells, with lambda phosphatase, reduced LRRK1 activity to levels observed in unstimulated cells (Fig 5C). This effect was largely prevented by using lambda phosphatase that had been preincubated with EDTA to suppress phosphatase activity (lambda phosphatase is a magnesium dependent phosphatase [42]) (Fig 5C).

### PKCα phosphorylates LRRK1 at 3 conserved residues lying in the GTPase COR_B_ domain

Kinase inactive LRRK1[20-2015, D1409A] was next phosphorylated in *vitro* in the presence or absence of PKCα, subjected to SDS-polyacrylamide gel electrophoresis, and digested with Trypsin + Lys-C (Fig 6A). The resultant peptides were analyzed either by Electron Activated Dissociation (EAD) on a ZenoTOF 7600 mass spectrometer or Higher energy Collisional Dissociation (HCD) on an orbitrap instrument, without phospho-peptide enrichment (Fig 6B & SFig 5). These analyses revealed a cluster of four phosphorylation sites regulated by PKCα phosphorylation, located within a loop of the COR_B_ domain (Thr1061, Ser1064, Ser1074 and Thr1075) (Fig 6B). The Thr1061 site was identified using both EAD and HCD fragmentation, Ser1064 was detected by the HCD method and the dual phosphorylated peptide encompassing Ser1074 and Thr1075 was detected using EAD method (Fig 6B & SFig4). Conservation evolutionary analysis employing the Consurf motif platform demonstrated that Ser1064 (score 9), Ser1074 (score 9) and Thr1075 (score 9) are highly conserved, but Thr1061 (score 1) is not. Significantly, Thr1075 lies within the optimal PKC consensus phosphorylation motif comprising basic Arg residues at the –3 and –5 positions in addition to a hydrophobic Phe residue at the +1 position [29] and these residues also display the highest conservation score of 9 (Fig 6C). Interestingly, these residues are located within a motif that is not conserved in LRRK2 (Fig 6D). We also used the HCD method to analyze phosphosites on LRRK1[20-2015, D1409A] ± phosphorylation with PKCα employing 2 additional protease digests (Asp-N and chymotrypsin) (SFig 5). This led to the identification of additional sites, several not induced by PKCα phosphorylation, that are likely phosphorylated by other kinases in insect cells.

**Figure 6.**
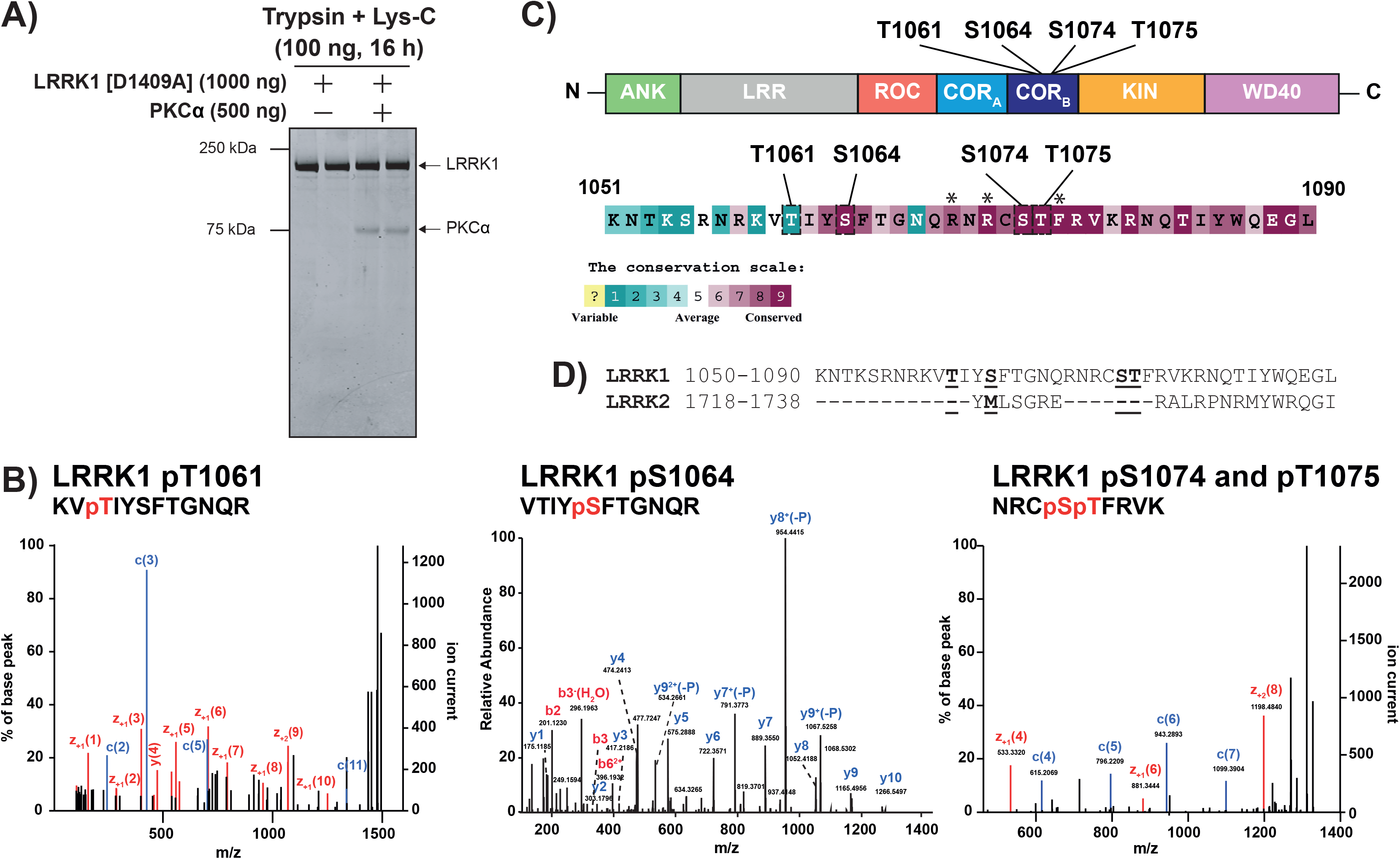
Ser1064, Ser1074 and Thr1075 are identified w/v spectrometry as three PKC-regulated sites on LRRK1. **(A)** Kinase inactive LRRK1[D1409A, 20-2015] (200 nM) was incubated ± PKCα (400 nM) the presence of MgATP. Reactions were terminated after 30 min with SDS-sample buffer and reactions resolved by SDS-polyacrylamide electrophoresis and gel stained by Coomassie blue. **(B)** The gel bands containing LRRK1 were digested with mixture of trypsin and Lys-C and the resultant peptides analyzed by Electron Activated Dissociation on a ZenoTOF 7600 mass spectrometer without phospho-peptide enrichment. The annotated MS/MS spectra of the three key PKC phosphorylated phosphosites encompassing phosphorylated Thr1061, phosphorylated Ser1064 and doubly phosphorylated Ser1074 and Thr1075 are shown. These sites were only detected in the LRRK1 samples phosphorylated with PKCα. Annotated MS/MS spectra of key PKC regulated phosphosites. MS/MS spectra of pT1061 and pS1064 were identified from HCD fragmentation acquired on Orbitrap Exploris 240 MS platform and the coverage of b and y ion series are denoted by red and blue color text respectively. MS/MS spectrum of a dual phosphorylated peptide of pS1074 and pT1075 sites were identified from EAD fragmentation acquired on Sciex Zeno-TOF 7600 platform. The EAD fragmentation enables accurate site localization and the coverage for c and z ion series were denoted with blue and red color text. **(C)** Upper panel displays the location within the COR_B_ domain arrangement of the Thr1061, Ser1064, Ser1074 and Thr1075 PKCα phosphorylation sites on the domain structure of LRRK1. Lower panels show the amino acid sequences encompassing the PKCα phosphorylation sites and their evolutionary conservation score from 1 (least conserved) to 9 (most conserved) of each residue determined calculated the Consurf server ((https://consurf.tau.ac.il/) [52]. Sequences of LRRK1 homologs were obtained from OrthoDB (https://www.orthodb.org/) [53] and alignment of sequences was performed using the MAFFT server (https://www.ebi.ac.uk/Tools/msa/mafft/) [54]. Subsequently, alignment is inserted into the Consurf server, with the sequence for human LRRK1 indicated as the query sequence (Taxonomy code 9606) and conservation scores are determined. Lower panel displays the conservation score scales utilized by the Cosurf server. Residues lying at the −5, −3 and +1 position of the Thr1075 phosphorylation site that confer optimal specificity for PKC isoform phosphorylation and marked with an asterisks **(D)** Sequence alignment of LRRK1 and LRRK2 surrounding the LRRK1 PKC phosphorylation sites showing that amino acid sequence across this region is not conserved between LRRK1 and LRRK2.

### PKC activation of LRRK1 is mediated by phosphorylation of Ser1064, Ser1074 and Thr1075

To explore the importance of the cluster of Thr1061, Ser1064, Ser1074 and Thr1075 PKC phosphorylation sites, we mutated these individually or in combination to Ala, to block phosphorylation (Fig 7A) or to Glu to mimic phosphorylation (Fig 7B). We found that mutation of Thr1075 to Ala alone, essentially blocked LRRK1 activation, whereas individual mutation of the other Ser1064 or Ser1074 only modestly impacted activity. Mutation of Thr1061 that has a low evolutionary conservation score suggesting that it would not play a critical role, had no effect (Fig 7A). The basal activity of the individual Ser1064Ala, Ser1074Ala and Thr1075Ala mutants were similar or marginally lower than that of wild type LRRK1 and significantly higher than that observed with kinase inactive LRRK1 (Fig 7A). Combined mutation of Ser1064Ala, Ser1074Ala and Thr1075Ala, reduced LRRK1 basal activity significantly and ablated phorbol ester induced activation (Fig 7A).

**Figure 7.**
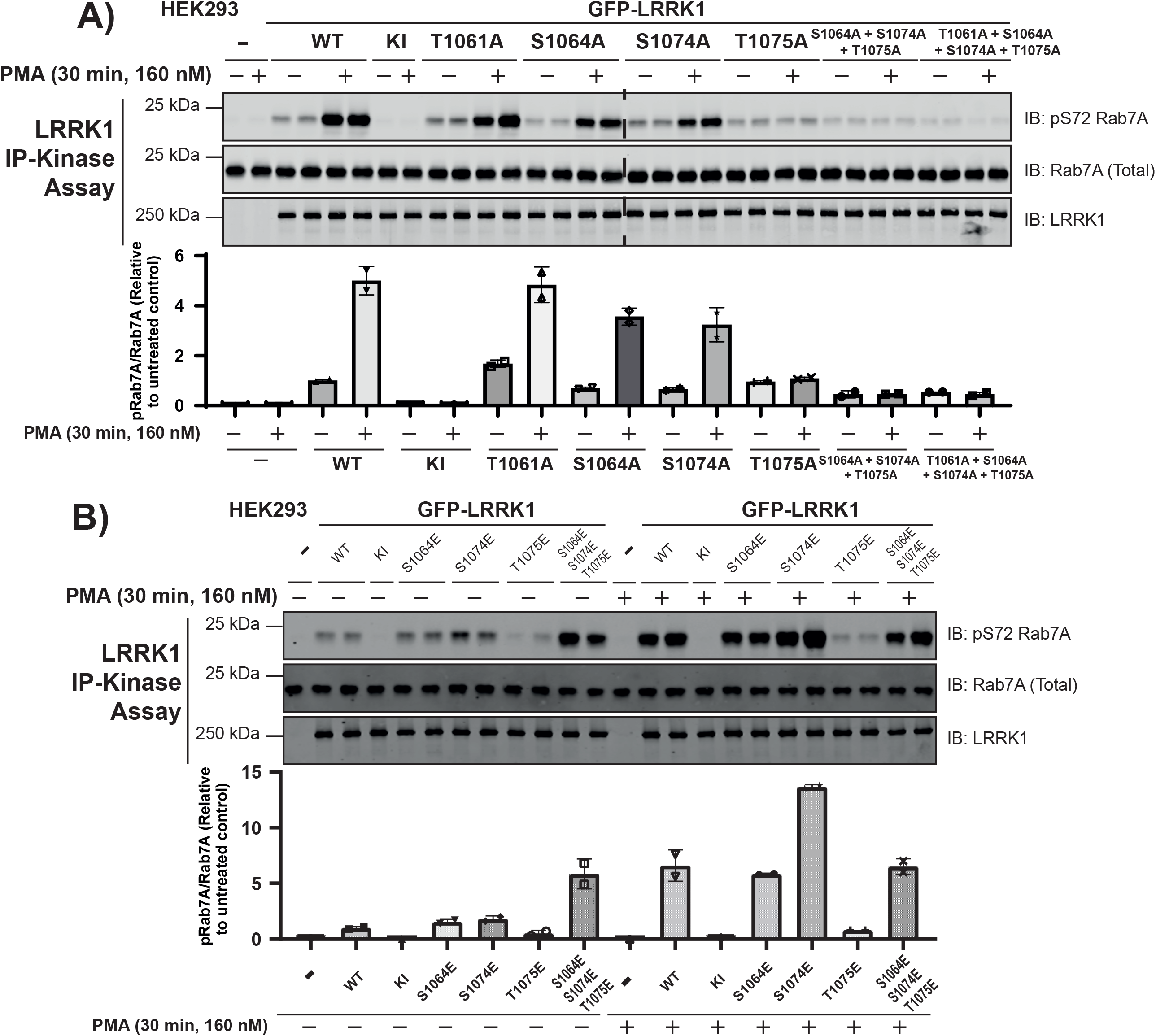
Mutations to Ser1064, Ser1074 and Thr1075 impact phorbol ester dependent LRRK1 activation. **(A and B)** HEK293 cells were transiently transfected with plasmids encoding wild type (WT), kinase-inactive (KI) LRRK1[D1409A] or the indicated mutants of GFP-LRRK1. Ala mutants to block phosphorylation were studied in (**A**) and Glu mutants to mimic phosphorylation were investigated in (**B**). 24 h post transfection, cells were serum-starved for 16 h and stimulated ± 160 nM phorbol 12-myristate 13-acetate (PMA) for 30 min. Cells were lysed and GFP-LRRK1 immunoprecipitated using an anti-GFP nanobody from 1 mg of cell extract. LRRK1 kinase activity towards recombinant Rab7A was assessed in a 30 min assay. Reactions were terminated by addition of SDS-sample buffer and levels of pSer72-Rab7A, Rab7A and LRRK1 were assessed by quantitative immunoblot analysis using the LI-COR Odyssey CLx Western Blot imaging system with the indicated antibodies. Combined immunoblotting data from 2 independent biological replicates (each performed in duplicate) are shown. Lower panel quantified immunoblotting data are presented as ratios of pRab7ASer72/total Rab7A (mean ± SEM) relative to levels observed with the activity of non-stimulated wild type LRRK1.

A triple Ser1064Glu, Ser1074Glu and Thr1075Glu mutant, displayed ~3-fold increased basal activity and was not activated further by phorbol ester treatment. Individual mutation of Ser1064 to Glu had no impact on the basal or phorbol ester induced activation of LRRK1, though mutating Ser1074 to Glu induces almost doubled kinase activity compared to the wild type protein following phorbol ester treatment. We found that Thr1075 to Glu inhibited LRRK1 activity and blocked its activation by phorbol ester.

### LRRK1 co-localizes with PKCα at the plasma membrane following phorbol ester stimulation

To investigate the localization of LRRK1, PKCα and Rab7A, we performed subcellular fractionation, generating membrane and cytosol fractions from unstimulated and phorbol ester stimulated cells. This revealed that as expected PKCα was mainly in the cytosol fraction in unstimulated cells and translocated to the membrane following phorbol ester stimulation (Fig 8A). LRRK1 was located both in the cytosol and membrane fractions and its distribution was not impacted by phorbol ester treatment. Most Rab7A and essentially all pSer72Rab7A were only observed in the membrane fraction (Fig 8A). The cytosolic fraction of total Rab7A is likely to represent the portion of the protein that is bound to GDI in cytosol. Immunofluorescence studies employing rapid immersion of cover slips in liquid nitrogen to extrude cytosolic proteins and enrich for membrane-associated proteins, confirmed that a significant proportion of LRRK1 was localized at the plasma membrane and that phorbol ester induced recruitment of PKCα to the plasma membrane, resulting in co-localization with the plasma membrane marker Na-K ATPase (Fig 8B, SFig 6). We observed a robust increase in pSer72 Rab7A staining with phorbol ester stimulation, though no obvious co-localization with LRRK1 is observed (Fig 8C). This study indicates that the pRab7A monoclonal antibody is suitable for immunofluorescence studies. We also observed that mutation of Ser1064, Ser1074 and Thr1075 to either Ala or Glu did not markedly influence the localization of LRRK1 observed (SFig 7).

**Figure 8.**
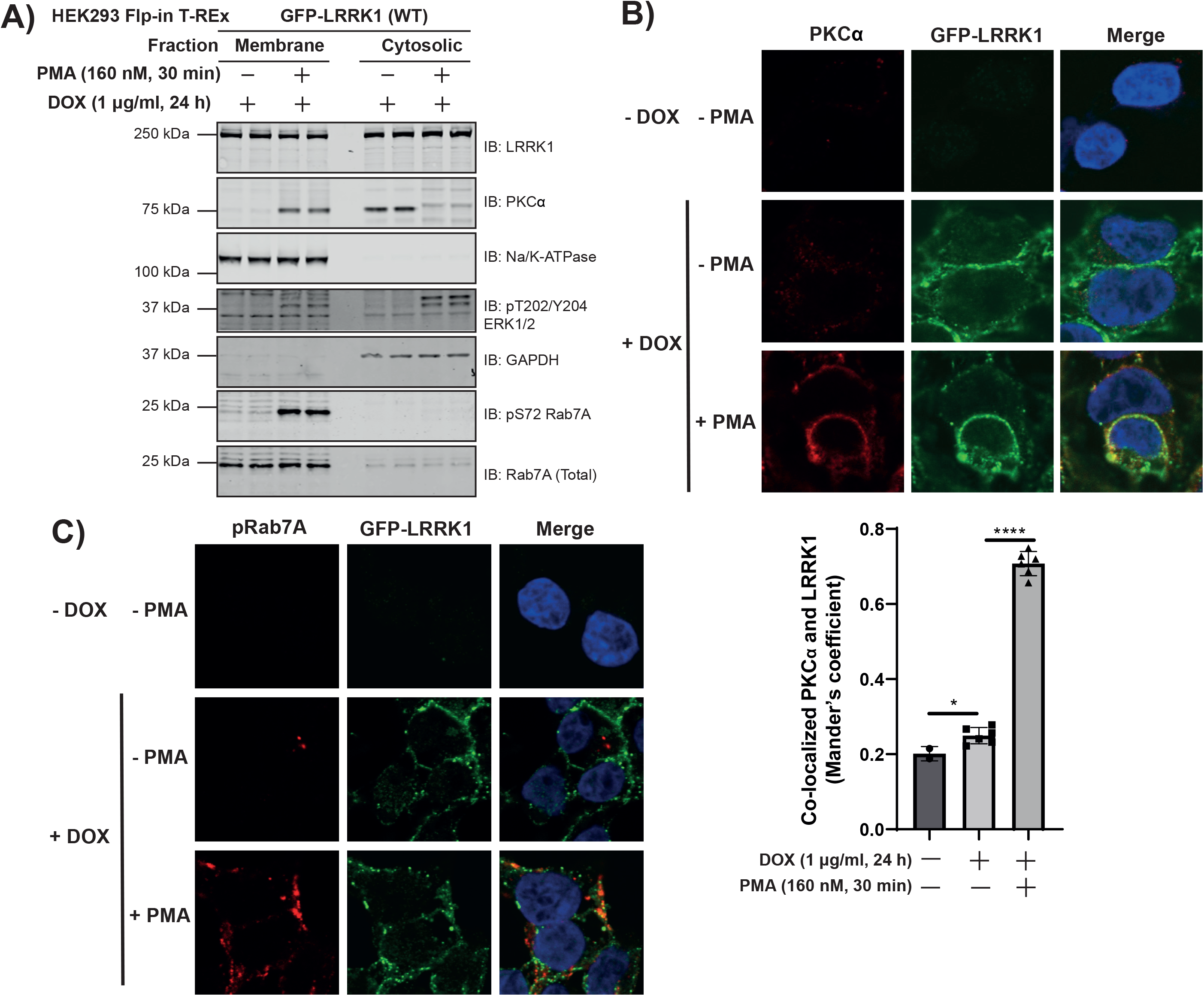
Phorbol ester induces co-localization of PKCα and LRRK1 at the plasma membrane. HEK293 Flp-In T-REx cells stably expressing wild type (WT) GFP-LRRK1 were treated with 1 μg/ml doxycycline for 24 h to induce GFP-LRRK1 expression. Cells were serum-starved for 16 h, and stimulated ± 160 nM phorbol 12-myristate 13-acetate (PMA) for 30 min. Next, cells were resuspended in a hypotonic solution (500 μl) and homogenized by passing through a 25-G needle. Extracts were fractionated into membrane and cytosolic fractions through high-speed ultracentrifugation. Membrane fractions were resuspended to the same volume as extract used for ultracentrifugation and equal volumes (500 μl) were assessed by quantitative immunoblot analysis using the LI-COR Odyssey CLx Western Blot imaging system with the indicated antibodies. Combined immunoblotting data from 2 independent biological replicates (each performed in duplicate) are shown. (**B, top panel**) As in (A) except cells were grown on coverslips and following treatments they were permeabilized by liquid nitrogen freeze–thaw to deplete cytosol [48] and then fixed and stained with mouse anti-PKCα, chicken anti-GFP and DAPI. (**B, bottom panel**) Co-localization of GFP-LRRK1 and PKCα was determined from a Mander’s coefficient (presented as mean ± SEM) after automatic thresholding. \**P* < 0.05, \*\*\*\**P* < 0.0001 by Student’s unpaired, two-tailed t test. **(C)** As in **(B, upper panel)** except cells are stained with rabbit anti-pS72 Rab7A and chicken anti-GFP.

## Discussion

Our key finding is that PKC isoforms directly activate LRRK1 by phosphorylating a cluster of 3 highly conserved PKC phosphorylation sites located on a loop of the COR_B_ domain (Ser1064, Ser1074 and Thr1075). Our data are consistent with a model in which phosphorylation of Thr1075, that lies within an optimal PKC phosphorylation motif, promotes phosphorylation of Ser1064 and Thr1074 that do not lie within optimal PKC site motifs, and that all 3 phosphorylation sites contribute to activity. Our data suggest that the Thr1075Glu mutation alone without phosphorylation of the two nearby residues results in loss of LRRK1 activity. It is possible that the Glu mutation is unable to mimic phosphorylation and induces a conformational change that is detrimental to basal kinase activity. It is also possible that phosphorylation of Thr1075 without concomitant phosphorylation of Ser1064 and Ser1074 is insufficient to activate LRRK1. Interestingly mutation of the adjacent Ser1074 site to Glu did not impact basal activity significantly, but enhanced phorbol ester stimulated activity 2-fold above that of the wild type. Thus, the most active species of LRRK1 that we have tested is the LRRK1[S1074E] mutant stimulated with phorbol ester (Fig 7B). In our mass spectrometry studies, we only observed a doubly phosphorylated Ser1074 and Thr1075 peptide, which also suggests that both these sites are efficiently phosphorylated by PKC isoforms. In future work it would be interesting to explore how phosphorylation of Thr1075 might regulate phosphorylation of the Ser1064 and Ser1074 residues. The Ser1064 and Ser1074 residues are unlikely to represent autophosphorylation sites, as the PKC phosphorylation site peptide mapping studies we undertook were performed using kinase-inactive LRRK1[20-2015, D1409A]. We also identified several additional phospho-sites on LRRK1, some of which have a high evolutionary score and likely to be phosphorylated in insect cells (SFig 5); further work is required to study any roles that these sites might play. The Ser1064, Ser1074 and Thr1075 or the residues encompassing these are not conserved in LRRK2 (Fig 6D) which likely accounts for why LRRK2 is not activated by PKC phosphorylation.

Recent work has reported a 5.8 Å Cryo-EM structure of the monomeric ROC-COR-Kinase-WD40 (RCKW) fragment of LRRK1 in the inactive conformation [2]. Although COR_B_ localized close to the kinase domain, it was not possible to model how PKC phosphorylation of the COR_B_ would impact the structure of LRRK1. The aC-helix is a universally conserved helix which is positioned centrally, between the N and C lobes of the kinase domain. The orientation of the helix is critical in kinase regulation with the distance between the N-terminal of the aC-helix and the activation loop defining the open and closed confirmations [43]. The AlphaFold [44] predicted structure of LRRK1 (https://alphafold.ebi.ac.uk/entry/Q38SD2), appears to be in the active conformation with a well ordered aC-helix. However, the region containing the Ser1064, Ser1074 and Thr1075 PKC phosphorylation sites is disordered which makes it difficult to predict how phosphorylation the COR_B_ loop impacts folding and interaction with the kinase aC-helix (Fig 9). We speculate that phosphorylation of LRRK1 promote ordering of residues in this region triggering stabilization of the aC-helix through an unknown mechanism. The current version of AlphaFold does not predict how phosphorylation impacts the structure. Recently, a 3.5 Å structure of an active tetrameric form of LRRK2 has been reported [18]. This consists of two peripheral LRRK2 molecules in the inactive conformation and two central LRRK2 protomers in an active conformation [18]. The residues in LRRK2 that lie at the equivalent region to Ser1064, Ser1074 and Thr1075 in the active LRRK2 subunits encompass residues 1721-1725 in LRRK2 (Fig 9, Fig 6C). Interestingly, in the active LRRK2 conformation, the 1721-1725 residues point towards the kinase domain and interact directly with the kinase αC-helix [18]. Furthermore, in a recent hydrogen-deuterium exchange mass spectrometry (HDX-MS) study, when the Type I inhibitor MLi-2 was added to induce the active conformation of LRRK2, stabilization of these 1721-1725 COR_B_ residues was observed [45]. The prediction from these observations was that phosphorylation of Ser1064, Ser1074 and Thr1075 could activate LRRK1 by engaging crosstalk between the C-terminal/aC-helix and Dk-helix, which is observed in the active state of LRRK2 (Fig 9). It would also be interesting to compare the subunit structures of the inactive and PKC phosphorylated active LRRK1. The finding that LRRK1 activity was inhibited by Triton-X100 could be explained if this disrupted multimeric subunit assembly required for activation (Fig 2A).

**Figure 9:**
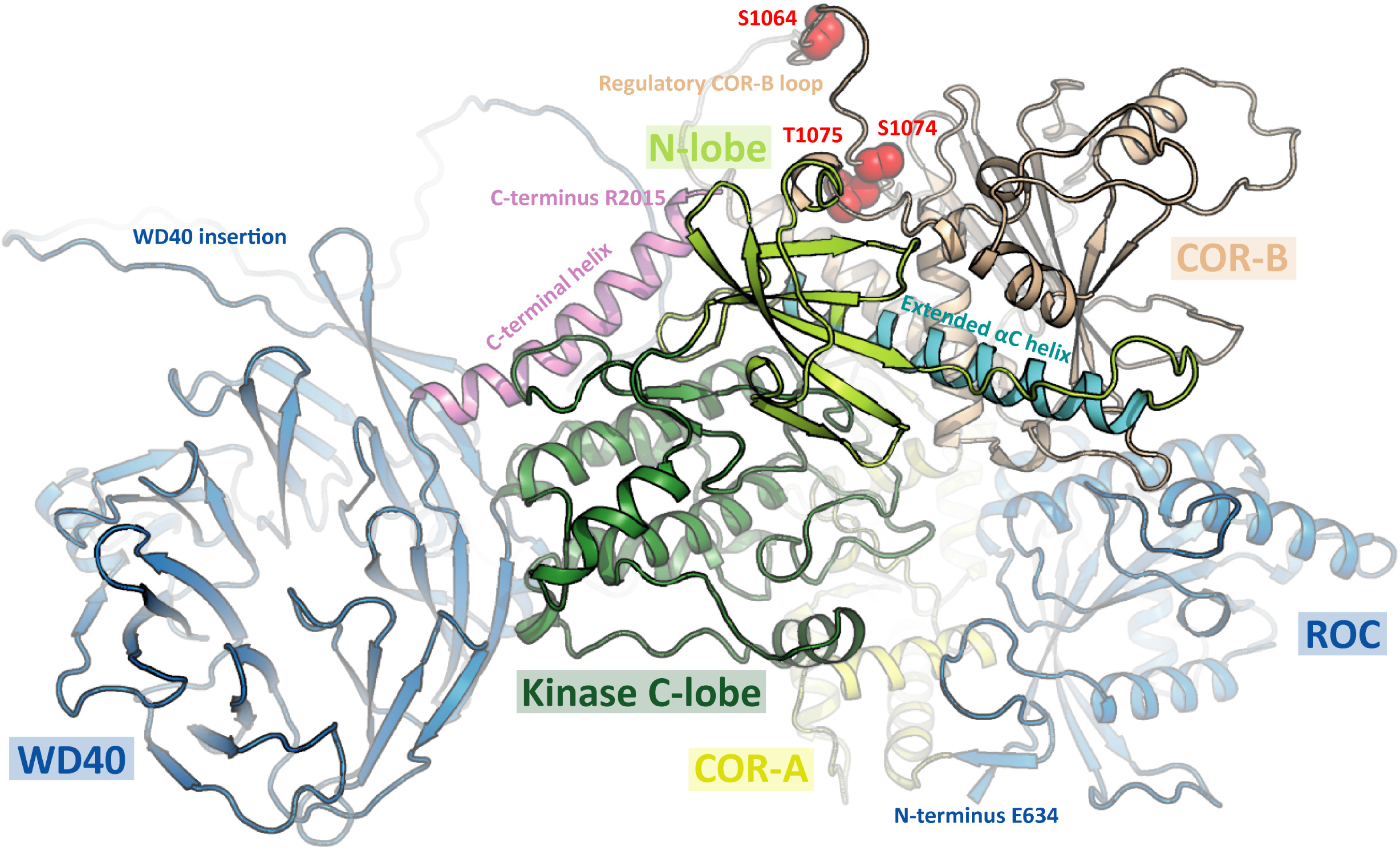
Model of LRRK1 structure. Prediction of LRRK1 (Q38SD2) structure as determined by AlphaFold Protein Structure Database (https://alphafold.ebi.ac.uk/entry/Q38SD2). Location of phosphosites (Ser1064, Ser1074 and Thr1075) within a disordered region are highlighted in red. The N-terminal ANK and LRR domains were omitted for clarity.

Based on our *in vitro* analysis all conventional and novel PKC isoforms that were tested activated LRRK1 to a similar level, suggesting that LRRK1 has evolved to be activated by multiple PKC isoforms. We observed that relatively low stoichiometry of PKC phosphorylation of ~0.23 mol ^32^P/mol protein was sufficient to activate LRRK1 ~100-fold (Fig 4B & SFig 2). Allowing for 3 PKC phosphorylation sites, theoretical maximal stoichiometry of phosphorylation of LRRK1 would be 3. Therefore, PKC phosphorylation can be estimated to enhance LRRK1 activity ~1000-fold emphasizing the major effect that phosphorylation has. The reasons for the relatively low phosphorylation levels of recombinant LRRK1 observed could be due the lack of enzyme and/or factors such as scaffolding proteins and/or membranes not being present in our in vitro reaction. After PKD, LRRK1 is to our knowledge the second protein kinase to be directly activated by PKC isoforms. It would be interesting to explore whether any previously described biology controlled by PKC isoforms is mediated through activation of LRRK1 and development of specific LRRK1 inhibitors would help address this question. It is also possible that other protein kinases including AGC kinases that are related to PKC isoforms and display similar substrate specificity, could also activate LRRK1 by phosphorylating Ser1064, Ser1074 and Thr1075. Further work would be required to study this and pinpoint which PKC isoform(s) mediate LRRK1 activation in vivo. It would also be important to generate phosphospecific antibodies that detect Ser1064, Ser1074 and Thr1075 phosphorylated activated form LRRK1 to better probe the function of the active form of this enzyme,

## Materials and Methods

### Reagents

[γ-^32^P]ATP was purchased from PerkinElmer. Phorbol 12-myristate 13-acetate (PMA) (#P8139), was purchased from Sigma-Aldrich. PKC-specific inhibitors GÖ6983 (#S2911), CRT0066051 (#S3422) and LXS-196 (#S6723) were purchased from Selleckchem. CHAPS hydrate (#C5070) and Digitonin (#D141) were purchased from Sigma-Aldrich. L-α-Phosphatidylserine (#840032C) and L-α-Diacylglyerol (#800815C) were purchased from Avanti Polar Lipids, Inc. Lambda Protein Phosphatase (#P0753S) was purchased from New England Biolabs. Trypsin/Lys-C mix (#V5073), Chymotrypsin (#V1062) and Asp-N (#V1621) were purchased from Promega. Bryostatin-1 (#2383) was purchased from TOCRIS. Recombinant Rab7A protein was expressed and purified as previously described [8]. Recombinant insect cell expressed PKC isoforms namely His-PKCα (DU5084), His-PKCβ (DU33630), Glutathione-S-transferase -PKCγ (DU30188) His-PKCε (DU33642), His-PKCθ (DU29920) and PKCζ (DU1447) were obtained from MRC Reagents and Services (https://mrcintranet.lifesci.dundee.ac.uk).

### Plasmids

All plasmids used in this study were obtained from the MRC PPU Reagents and Services (https://mrcppureagents.dundee.ac.uk). Each plasmid was confirmed by sequencing at the MRC Sequencing and Services (https://www.dnaseq.co.uk) and are available to request via the MRC PPU Reagents and Services website (https://mrcppureagents.dundee.ac.uk).

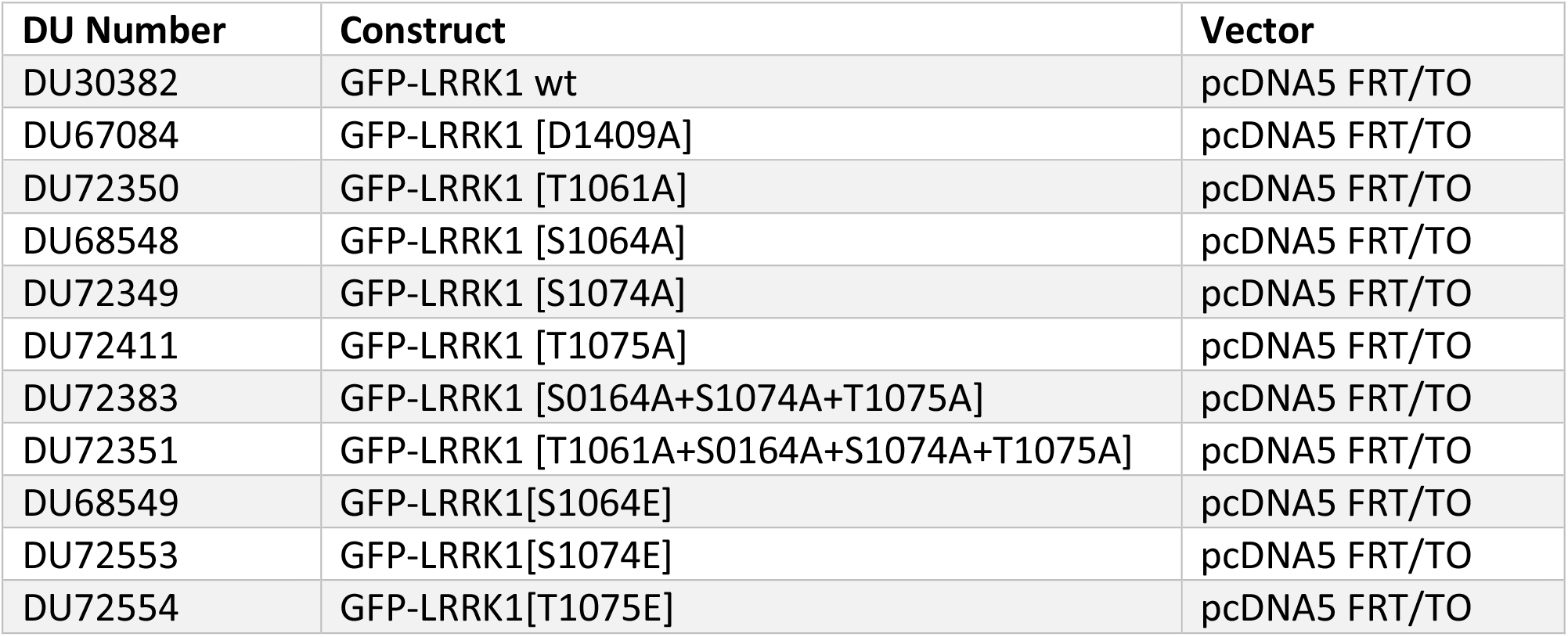
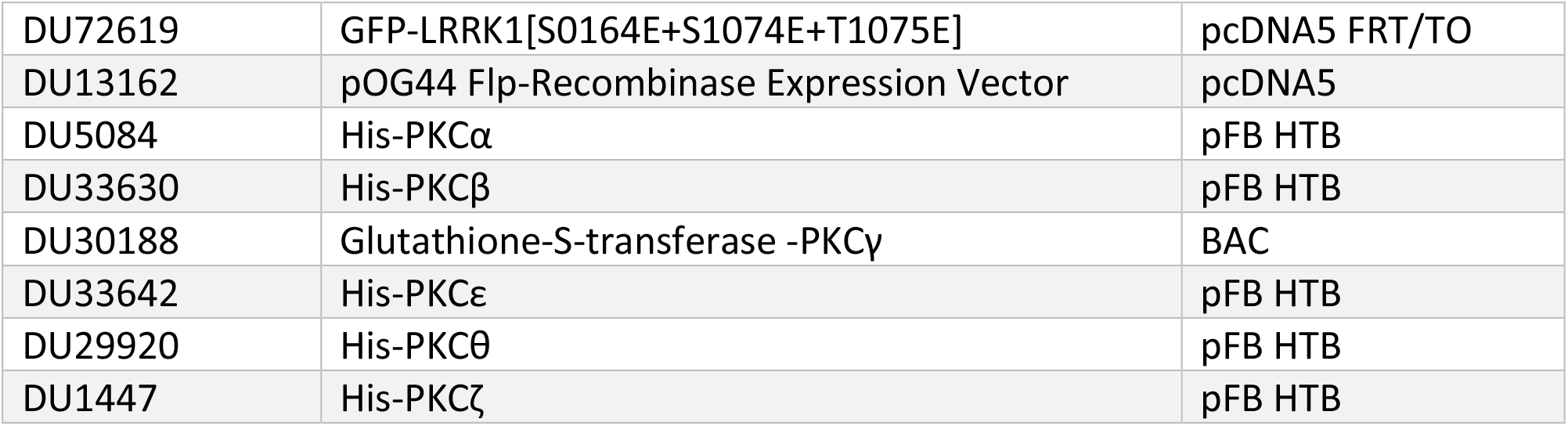

### Antibodies

All antibodies were diluted in 5% (w/v) BSA (bovine serum albumin) in TBST Buffer (20 mM Tris-HCl pH 7.5, 0.15 M NaCl and 0.1% (v/v) Tween-20) and used for immunoblotting analysis at the indicated concentration or dilution.

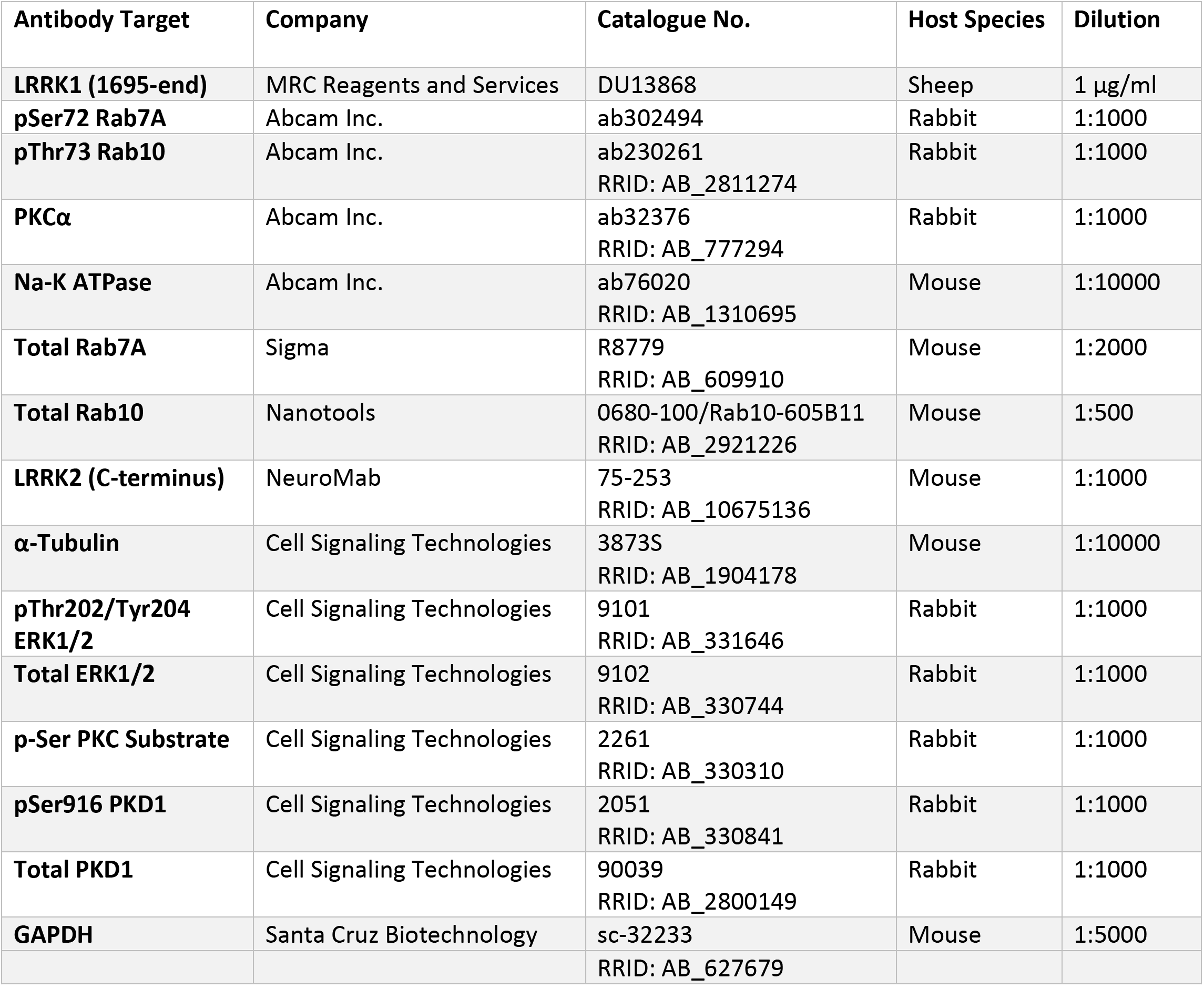

### LRRK1 KO primary MEF generation

Wild-type and homozygous LRRK1 knock-out mouse embryonic fibroblasts (MEFs) were isolated from littermate-matched mouse embryos at day E12.5 resulting from crosses between heterozygous wildtype/KO mice as described previously [8].

### Cell Culture and Lysis

HEK293 wild type (ATCC, #CRL-1573) and HEK293 Flp-in T-REx (ThermoFisher, #R78007) wild type cells were cultured in Dulbecco’s Modified Eagle’s Medium (DMEM; GIBCO, #11960-085) containing 10% (v/v) fetal bovine serum (Sigma, #F7524), 2 mM L-glutamine (GIBCO, #25030024), 100 U/ml penicillin-streptomycin (Invivogen, #15140122). HEK293 Flp-in T-REx wild type cells were grown in DMEM containing 10% (v/v) fetal bovine serum, 2 mM l-glutamine, 100 U/ml penicillin-streptomycin supplemented with 15 μg/ml Blasticidin (Invivogen, #ant-bl-10p) and 50 μg/ml Zeocin (Invivogen, #ant-zn-5). MEFs were grown in DMEM containing 10% (v/v) fetal bovine serum, 2 mM L-glutamine, 100 U/ml penicillinstreptomycin, supplemented with 1 mM sodium pyruvate (GIBCO, #11360-039) and nonessential amino acids (GIBCO, #11140-035). All cells were incubated at 37 °C, 5% CO_2_ (v/v) in a humidified atmosphere and regularly tested for mycoplasma contamination. Cells were lysed in an ice-cold lysis buffer containing 50 mM HEPES, pH 7.4, 0.3% CHAPS hydrate (v/v), 270 mM sucrose, 150 mM NaCl, 1 mM sodium orthovanadate, 50 mM NaF, 10 mM 2-glycerophosphate, 5 mM sodium pyrophosphate, 1 μg/ml microcystin-LR (Enzo Life Sciences, #ALX-350-012-M001) and complete EDTA-free protease inhibitor cocktail (Sigma, #11836170001). Lysates were clarified by centrifugation at 20,800 g at 4°C for 10 min and supernatant protein concentrations quantified using the Bradford assay kit (ThermoFisher, #23236)

### Generation of HEK293 Flp-in T-REx stable cell lines

Non-transfected HEK293 Flp-in T-REx wild type cells were seeded into 10 cm dishes at 4.5 x 10^6^ cells/well. Cells were transfected using Polyethylenimine method [47]. Briefly, 0.5 μg of LRRK1 or LRRK2 plasmids, 4.5 μg of pOG44 Flp-Recombinase Expression Vector (ThermoFisher, #V600520) and 15 μl of 1 mg/ml PEI were added to 1 ml OptiMEM reduced serum media (GIBCO, #31985-062) and vortexed for 20 seconds. Solution was then incubated at room temperature for 20 min to allow formation of DNA-PEI complexes. Transfection mixes were subsequently added to cell culture medium and allowed to incubate at 37 °C. 24 h post transfection, cells were selected for successful transfection using DMEM containing 10% (v/v) fetal bovine serum, 2 mM l-glutamine, 100 U/ml penicillin, and 100 μg/ml streptomycin supplemented with 15 μg/ml Blasticidin and 100 μg/ml Hygromycin (Invivogen, #ant-hg-5) for 48 hours. Cells were then allowed to recover from selection in Hygromycin by incubation in DMEM containing 10% (v/v) fetal bovine serum, 2 mM l-glutamine, 100 U/ml penicillinstreptomycin for 1 week.

### Transient and stable overexpression experiments and cell treatments

24 h prior to transfection or treatment, HEK293 cells were seeded into 10 cm dishes at 4.5 x 10^6^ cells/well. Cells were transfected using Polyethylenimine method [47], as described above.

Briefly, 5 μg of LRRK1 plasmids and 15 μl of 1 mg/ml PEI were added to 1 ml OptiMEM reduced serum media (GIBCO, #31985-062) and vortexed for 20 seconds. Solution was then incubated at room temperature for 20 min to allow formation of DNA-PEI complexes. Transfection mixes were subsequently added to cell culture medium and allowed to incubate at 37 °C. For stable overexpression, HEK293 Flp-in T-REx stable cell lines were induced to express GFP-LRRK1 or LRRK2 by treatment with 1 μg/ml Doxycycline (Sigma, #D9891) for 24 h. For experiments involving phorbol ester stimulation, cells were incubated in DMEM not containing serum overnight. Cells were then treated with 160 nM phorbol 12 -myristate 13-acetate (PMA; Sigma, #P8139) for the time indicated in the figure legends. PMA and kinase inhibitors GÖ6983, CRT0066051 and LXS-196 were all made up in DMSO at a 1000-fold higher concentration than used in cells, which were treated with the indicated doses of inhibitor at 1:1000 dilution for the indicated time. An equivalent volume of DMSO was added to cells not treated with an inhibitor or phorbol ester.

### Immunoblotting analysis

Lysates were mixed with a quarter of volume or 4x SDS-PAGE sample buffer [50 mM Tris–HCl, pH 6.8, 2% (w/v) SDS, 10% (v/v) glycerol, 0.02% (w/v) Bromophenol Blue and 1% (v/v) 2-mercaptoethanol] (Novex) and heated at 95 °C for 5 min. 10–20 μg samples were loaded on to 4-12% NuPAGE™ Bis-Tris gels (Invitrogen, #NP0322BOX) and electrophoresed at 90 V for 15 min, then 150 V for 75 min in NuPAGE™ MOPS buffer (Invitrogen, #NP0001). Proteins were transferred onto nitrocellulose membranes (GE Healthcare, Amersham Protran Supported 0.45 μm, #10600041), at 90 V for 90 min on ice in transfer buffer (48 mM Tris-HCl, 39 mM glycine; freshly supplemented with 20% methanol (v/v)). Transferred membranes were blocked with 5% (w/v) milk in TBST Buffer at room temperature for 60 min. Membranes were then incubated with primary antibodies diluted in blocking buffer overnight at 4 °C. After washing membranes in TBST, membranes were incubated at room temperature for 1 h with near-infrared fluorescent IRDye antibodies (LI-COR) diluted 1:10000 in TBST and developed using the LI-COR Odyssey CLx Western Blot imaging system and signal quantified using the Image Studio software. Gathered data was subsequently analysed and plotted in GraphPad Prism (version 9.1.0). The method used for immunoblotting analysis for the LRRK1 pathway further described in detail in a protocols.io method (dx.doi.org/10.17504/protocols.io.6qpvr68e3vmk/v1).

### Cloning, expression, and purification of LRRK1 wild type, kinase inactive [D1409A]

Briefly, the DNA coding for the human LRRK1 residues 20 to 2015 (OHu72031 from Genscript) was PCR-amplified using the forward primer TACTTCCAATCCGCTGTGTGTCCAGAACGTGCCATGG and the reverse primer TATCCACCTTTACTGTCACCTTCTCTTGCGAGTGCAAGCCTCC. The T4 polymerase-treated amplicon was inserted into the transfer vector pFB-6HZB (SGC) by ligation-independent cloning. The respective point mutations were introduced applying the QuikChange method. The resulting plasmids were utilized for the generation of recombinant Baculoviruses according to the Bac-to-Bac expression system protocol (Invitrogen #10359016). Exponentially growing Spodoptera frugiperda 9 cells (2E06 cells/mL in Lonza Insect-XPRESS medium #BELN12-730Q) were infected with high-titer Baculovirus suspension. After 66 h of incubation (27°C and 90 rpm), cells were harvested by centrifugation. The expressed protein construct contained an N terminal His6 Z tag, cleavable with TEV protease. For LRRK1 purification, the pelleted Spodoptera frugiperda 9 cells were washed with PBS, re-suspended in lysis buffer (50 mM HEPES pH 7.4, 500 mM NaCl, 20 mM imidazole, 0.5 mM TCEP, 5% (v/v) glycerol) and lysed by sonication. The lysate was cleared by centrifugation and loaded onto a Ni NTA column. After vigorous rinsing with lysis buffer the His6-Z tagged protein was eluted in lysis buffer containing 300 mM imidazole. Immediately thereafter, the eluate was diluted with buffer containing no NaCl, to reduce the NaCl-concentration to 250 mM and loaded onto an SP-Sepharose column. His6 Z TEV-LRRK1 was eluted with a 250 mM to 2.5 M NaCl gradient and treated with TEV protease overnight to cleave the His6 Z tag. Contaminating proteins, the cleaved tag, uncleaved protein and TEV protease were removed by another combined SP-Sepharose Ni NTA step. Finally, LRRK1 was concentrated and subjected to gel filtration in storage buffer (20 mM HEPES pH 7.4, 150 mM NaCl, 0.5 mM TCEP, 5% glycerol) using an AKTA Xpress system combined with an S200 gel filtration column. The final yield as calculated from UV absorbance was 0.1 mg/L. The expression method for LRRK1 is further described in a detailed protocols.io method (dx.doi.org/10.17504/protocols.io.b7ternje).

### Immunofluorescence

For analysis of LRRK1 co-localization with pSer72 Rab7A, PKCα or Na-K ATPase, a freeze-thaw protocol for cell fixation was employed which enables extrusion of cytosolic proteins and enrichment of membrane-associated proteins [48]. Briefly, HEK293 Flp-in T-REx/GFP-LRRK1 cells were seeded at 2.5 x 10^5^ cells/well in 6 well plates on glass coverslips (VWR, 631-0125; square 22×22 mm thickness 1.5). HEK293 Flp-in T-REx stable cell lines were induced to express GFP-LRRK1 by treatment with 1 μg/ml Doxycycline for 24 h. For investigation of how LRRK1 phosphosite mutations affect localization, HEK293 wild type cells were seeded at 2.5 x 10^5^ cells/well in 6 well plates on glass coverslips. HEK293 cells were then transiently transfected with the indicated GFP-LRRK1 plasmids for 24 h. Subsequently cells were incubated overnight in serum-free media overnight and then treated with 160 nM PMA for a duration of 30 min. Cells were next washed twice with ice-cold PBS buffer (137mM NaCl, 2.7mM KCl, 10mM Na_2_HPO_4_, 1.8 KH_2_PO_4_, pH 7.4) and twice with ice-cold glutamate lysis buffer (25mM HEPES, pH 7.4, 25 mM KCl, 2.5 mM magnesium acetate, 5 mM EGTA, 150 mM potassium glutamate). Coverslips were removed from wells with excess liquid removed by touching the edge on a paper towel. Coverslips were then snap frozen in liquid nitrogen and allowed to thaw at room temperature on a paper towel before replacement into the well. Cells were washed twice with ice-cold glutamate lysis buffer before fixing with 4% (v/v) paraformaldehyde at room temperature for 20 min. Next, unreacted paraformaldehyde was quenched by washing coverslips twice with DMEM (supplemented with 10 mM HEPES, pH 7.4) followed by blocking with 1% (w/v) BSA in PBS for 30 min. The coverslips were incubated with chicken polyclonal anti-GFP (#ab13970, Abcam, Inc.) primary antibody, diluted 1:1000 and rabbit monoclonal anti-PKCα (#ab32376, Abcam, Inc.) primary antibody, diluted 1:500 or mouse monoclonal anti-Na-K ATPase (#ab76020, Abcam, Inc.) primary antibody, diluted 1:500 or rabbit monoclonal anti-pSer72 Rab7A (#302494, Abcam, Inc.) in 1% (w/v) BSA in PBS buffer for 1 h at room temperature followed with 3 times for 15 min washes with 0.2% (w/v) BSA in PBS. Then the coverslips were incubated with Alexa Fluor^®^ secondary antibody (Invitrogen A-31553) diluted 1:500 in 1% (w/v) BSA in PBS for 1 h. The coverslips were washed 3 times for 15 min with 0.2% (w/v) BSA in PBS and rinsed with distilled water just before mounting. ProLong^®^ Gold Antifade Reagent with DAPI (ThermoFisher, #P36935) was used to mount the coverslips on glass slides (VWR, 631-0117). Slides were then imaged using Leica TCS SP8 MP Multiphoton Microscope using a 40x oil immersion lens choosing the optimal imaging resolution with 1-pixel size of 63.3 nm x 63.3 nm. Images were subsequently analysed using Fiji (https://fiji.sc/) and colocalization was quantified using the Fiji plug-in, JaCoP (https://imagej.net/plugins/jacop) which was used to determined Mander’s coefficients using automatic thresholds to determine overlap in signal between channels. Coefficients were then statistically analysed in GraphPad Prism (version 9.1.0) using Student’s unpaired, two-tailed t tests. This immunofluorescence method is further described in detail in a protocols.io method (dx.doi.org/10.17504/protocols.io.ewov1nmzkgr2/v1)

### Membrane Fractionation

For crude membrane fractionation, a previously described method was employed [14]. Briefly, 15 cm dishes of HEK293 Flp-in T-REx/GFP-LRRK1 cells were seeded at 1.8 x 10^7^ cells/dish. Cells were then induced to express GFP-LRRK1 by treatment with 1 μg/ml Doxycycline for 24 h. Subsequently cells were incubated overnight in serum-free media and then treated with 160 nM PMA for a duration of 30 min. Cells were washed twice with ice-cold phosphate buffered saline (PBS; 137 mM NaCl, 2.7 mM KCl, 10 mM Na_2_HPO_4_ and 1.8 mM KH_2_PO_4_) resuspended in 400 μl of a hypotonic solution (10 mM HEPES pH 7.4, 50mM NaF, 5 mM Sodium Pyrophosphate and complete EDTA-free protease inhibitor cocktail (Roche). After 15 min of incubation in hypotonic solution, 100 μl of resuspension buffer (250 mM HEPES, pH 7.4, 750 mM NaCl, 5 mM MgCl_2_, 2.5 mM DTT, 500 nM GDP, 250 mM NaF, 25 mM Sodium Pyrophosphate, 1 μg/ml microcystin-LR and complete EDTA-free protease inhibitor cocktail (Roche)) was added. Cells were homogenized and lysed by passing through a 25-G needle 25 times. Lysates were then centrifuged at 2,000 g for 5 min at 4 °C to pellet nuclei and mitochondrial fraction and the supernatant containing membrane fraction and cytosol was collected. The membrane fraction was then pelleted by ultracentrifugation at 100,000 g for 25 min at 4 °C. Supernatant which contained the cytoplasmic fraction was collected and carefully transferred into a new tube without disturbing the pellet fraction. Membrane fraction pellet was then washed twice gently with 500 μl is ice cold PBS buffer to remove any cytoplasmic contaminants before being resolubilized in 500 μl lysis buffer (50 mM HEPES, pH 7.4, 1% (v/v) Triton X-100, 150 mM NaCl, 1 mM MgCl_2_, 0.5 mM DTT, 100nM GDP, 50mM NaF, 5 mM Sodium Pyrophosphate, 1 μg/ml microcystin-LR and complete EDTA-free protease inhibitor cocktail (Roche)). Resolubilized membrane fractions were then centrifuged at 2,000 g for 5 minutes at 4 °C with resultant supernatant containing the solubilized membrane fraction collected into a new tube. The protocol for the membrane fractionation method is further described in detail in a protocols.io method (dx.doi.org/10.17504/protocols.io.yxmvmnb99g3p/v1).

### GFP-LRRK1 Immunoprecipitation for kinase assay

Briefly, cells were harvested in lysis buffer and clarified by centrifugation at 11,000 g for 10 min at 4 °C. 1000 μg of whole cell lysate was incubated with 10 μl of packed nanobody anti-GFP binder-Sepharose beads (generated by the MRC PPU Reagents and Services) for 2 h. Bound complexes were recovered by washing the beads twice with 25 mM HEPES pH 7.5, 500 mM NaCl, 10 mM GTP-γ-S and once with 25 mM HEPES pH 7.5, 150 mM NaCl, 10 mM GTP-γ-S. Bound complexes were then equilibrated in HEPES kinase buffer (25 mM HEPES, 50 mM KCl and 0.1 % (v/v) 2-mercaptoethanol) by washing once. The beads were then directly subjected to a Rab7A kinase assay by incubation in 20 μl of a mixture containing (1 μM Rab7A, 1 mM nonradioactive ATP, 10 mM MgAC, 25 mM HEPES, 50 mM KCl and 0.1 % (v/v) 2-mercaptoethanol). The reactions were incubated at 30 °C for 30 min and terminated by the addition of 20 μl of 2X SDS Sample Buffer containing 1% (v/v) 2-mercaptoethanol. The samples were heated for 10 min at 97 °C and the beads separated from the reaction by centrifugation though 0.22 μm Spin-X columns (CLS8161, Sigma) for 5 min at 3, 500 g. 10 μl of the resultant sample was analyzed by quantitative immunoblot analysis as described above. The immunoprecipitation kinase assay for GFP-LRRK1 is described in further detail in a protocols.io method (dx.doi.org/10.17504/protocols.io.kqdg3p2wpl25/v1).

### Immunoprecipitation and kinase activity for endogenous LRRK1

0.5 mg of affinity purified sheep polyclonal total LRRK1 antibody (Sheep number S405C, DU13868) at a concentration of 0.23 mg/ml in PBS buffer was covalently coupled to 25 mg of Dynabeads™ M-270 Epoxy magnetic beads (ThermoFisher, #14302D), according to the manufacturer’s protocol. Briefly, the 25 mg of magnetic beads were resuspended in 0.24 ml of C1 Buffer (0.1 M Sodium Phosphate (Na_2_HPO_4_:NaH_2_PO_4_) pH 7.4). 2.4 ml of C2 Buffer (3 M (NH_4_)_2_SO_4_) in 0.1 M Sodium Phosphate pH 7.4) was added. The resulting mixture added to 2.16 ml of the 0.23 mg/ml total LRRK1 antibody and incubated for 16 h at 37 °C. The antibody-bead complex was isolated using a magnet separator (DynaMag™-2, Life Technologies #12321D) and washed in 1 ml of buffer HB (100 mM Glycine pH 11.3), vortexed for 5 sec and immediately washed again in 1 ml of LB Buffer (200 mM Glycine pH 2.8) with 5 sec vortexing. This was followed by two washes in 1 ml of SB Buffer (50 mM Tris-HCl pH 7.4, 140 mM NaCl, 0.1% (v/v) Tween-20). The beads were resuspended in 1 ml of SB Buffer and incubated on a roller mixer at room temperature for 10 min. Finally, the coupled antibody-bead complex was isolated and resuspended to give a final concentration of 10 mg of antibody-bead/ml.

For the immunoprecipitations, the equivalent of 2 mg of extracts derived from unstimulated and phorbol ester stimulated wild type and LRRK1 knock-out cells prepared in 0.3% (w/v) CHAPS lysis buffer, were incubated with 50 μl of the 10 mg/ml antibody-bead complex for 2 h on a shaking platform at 4 °C. The antibody-bead complexes were washed twice with 25 mM HEPES pH 7.5, 500 mM NaCl, 10 μM GTP-γ-S, once with 25 mM HEPES pH 7.5, 150 mM NaCl, 10 μM GTP-γ-S and finally once in kinase assay buffer (25 mM HEPES, 50 mM KCl and 0.1 % (v/v) 2-mercaptoethanol). The beads were subjected to a Rab7A kinase assay as described above. The samples are heated for 10 min at 97 °C and removed using a magnet. 10 μl of the resultant sample was analysed by quantitative immunoblot analysis as described above.

### Phosphorylation and activation of LRRK1 with PKC isoforms

PKC activation kinase assays were conducted in two distinct steps. In the first step, indicated PKC isoforms (100 nM) were incubated with either GFP-LRRK1 immunoprecipitated from 1 mg of HEK293 Flp-in T-REx/GFP-LRRK1 wild type cell extract or with recombinant LRRK1 (50 nM, residues 20-2015) in a 20 ml final reaction mixture containing 50 mM HEPES pH 7.5, 0.1 % (v/v) 2-mercaptoethanol, 100 μg/ml L-α-Phosphatidylserine, 10 μg/ml L-α-Diacylglyerol, 1 mM CaCl_2_ and 10 μM GTP-γ-S, 10 mM MgAc and either 1 mM non-radioactive ATP or where indicated 0.1 mM [γ-^32^P]ATP. The reaction mixture was incubated for 30 min at 30 °C. In a secondary step, reactions were supplemented with 10 μl of a master mix containing 1 mM Rab7A and either 1 mM non-radioactive ATP or when indicated, 0.1 mM [γ-^32^P]ATP. The second step of the kinase reaction was carried out at 30 °C for the timepoints indicated in the figure legend and reactions terminated by the addition of 10 μl of 4x SDS–PAGE sample buffer containing 1% (v/v) 2-mercaptoethanol. 25% (10 μl) of total reaction volume was used for each immunoblot analysis. For reactions where [γ-^32^P]ATP was employed, 75% (30 μl) of total reaction volume was analyzed by autoradiography. Briefly, proteins were resolved by SDS-PAGE and detected with Coomassie staining. The gels were imaged with an EPSON scanner, then sandwiched and secured between two sheets of pre-wet cellophane (Bio-Rad) and subsequently dried in a GelAir dryer for 45–60 min. Dried gels were exposed to Amersham Hyperfilm MP overnight (VWR, #28-9068-42), in an autoradiography cassette and the films were later developed using a Konica auto-developer. The method used for activation of LRRK1 by PKC phosphorylation analysis is described in further detail in a protocols.io method (dx.doi.org/10.17504/protocols.io.5jyl89d5rv2w/v1).

### Determining Stoichiometry of PKCα-mediated LRRK1 phosphorylation

LRRK1 was subjected to PKCα phosphorylation using 0 to 100 nM PKCα as described above using [γ-^32^P]ATP of specific activity of ~500 cpm/pmol. Reactions were undertaken in quadruplicate. After polyacrylamide gel electrophoresis the dried Coomassie band corresponding to LRRK1 was excised and ^32^P content quantified by Cherenkov counting using liquid scintillation analyzer (TriCarb 4910TR, Perkin Elmer). The ratio of ^32^P radioactivity/mol of LRRK1 was calculated.

### *in vitro* λ (Lambda) phosphatase treatments

Following immunoprecipitation of GFP-LRRK1 (immunoprecipitated from 1 mg of HEK293 Flp-in T-REx/GFP-LRRK1 WT cell extract), immunoprecipitates are washed once with 500 μl of 1X phosphatase buffer (New England Biolabs, #B7061SVIAL). Washed immunoprecipitates were then incubated with 200U of λ phosphatase (0.5 μl of stock 400,000 U/ml) for 30 min at 30 °C in a reaction containing 1X phosphatase buffer supplemented with 10 mM MnCl_2_. For the inactivated phosphatase controls, λ phosphatase activity was inhibited by pre-incubation of λ phosphatase with 50 mM EDTA for 10 min to enable complete inhibition prior to addition to immunoprecipitates.

### Phosphosite identification by Mass Spectrometry

A 20 μl reaction mixture was set up containing 200 nM of kinase inactive LRRK1[D1409A, 20-2015] (1 μg) containing 400 nM recombinant full-length PKCα (500 ng) and in a buffer containing 50 mM HEPES pH 7.5, 0.1 % (v/v) 2-mercaptoethanol, 100 μg/ml L-α-Phosphatidylserine, 10 μg/ml L-α-Diacylglyerol, 1 mM CaCl_2_ and 10 mM GTP-γ-S, 10 mM MgAc and 1 mM ATP (non-radioactive) for 30 min at 30 °C. The reactions were stopped by the addition of 6.6 ml of 4× SDS–PAGE sample buffer containing 1% (v/v) 2-mercaptoethanol, and samples subjected to electrophoresis on a 4–12% SDS–PAGE gels. The gel was stained with Coomassie blue and the bands corresponding to LRRK1 were excised and destained by three 10 min washes in 40% (v/v) acetonitrile in 40mM NH_4_HCO_3_. Gel pieces were then reduced by incubation in 5 mM DTT in 40mM NH_4_HCO_3_ at 56°C for 30 min. The DTT solution was removed by careful pipetting and washed in 40% (v/v) acetonitrile in 40 mM NH_4_HCO_3_ for 10 min. This was followed by alkylation of the samples by addition of 20 mM iodoacetamide in 40 mM NH_4_HCO_3_ for 20 min at room temperature. Gel pieces were subsequently dehydrated by washing in 100% (v/v) acetonitrile for 10 min. Acetonitrile is removed by careful pipetting and vacuum drying for 10 min to remove any residual acetonitrile. Proteases (w/v) were freshly prepared in the appropriate buffers (Trypsin + LysC in 50 mM TEABC, Asp-N in 50 mM Tris-HCl, Chymotrypsin in 100 mM Tris-HCl + 10 mM CaCl_2_) and 100 ng of proteases were added to gel pieces and incubated overnight on a thermomixer at 37°C, 800 xg. Digested peptides were extracted from gel pieces by addition of 200 μl of extraction buffer (80% (v/v) acetonitrile in 0.2% (v/v) formic Acid), incubated on a Thermomixer with an agitation at 800 xg for 20 min. The extraction was completed by repeating this step again. The eluate was immediately vacuum dried and resuspended in 80 μl of Solution A (0.1% (v/v) TFA (trifluoroacetic acid). Resuspended peptides were then subjected to a C18 clean-up using in-house prepared C18 stage tips. One C18 disk was prepared by punching with 16-gauge needle and loaded on to 250 μl pipette tips. C18 resin was activated by addition of 100 μl of acetonitrile and centrifuged for 2 min at 2,000 xg. Flow through was discarded and C18 resin was equilibrated by addition of 80 μl of Solution A (0.1% (v/v) TFA) and centrifuged for 2 min at 2,000 xg. Flow through were discarded again and peptide digests were loaded onto C18 resin and centrifuged at 1,500 xg for 5 min. The flow through was re-applied and subsequently centrifuged at 1,500g for 5 min. C18 resins were then washed twice by addition of 80 μl of Solution A (0.1% (v/v) TFA) and centrifuged for 2 min at 2,000 xg. Peptides were eluted by addition of 80 μl of Solution B (40% (v/v) acetonitrile in 0.1% (v/v) TFA) and centrifuged for 2 min at 1,500 xg. Eluates were immediately snap frozen and vacuum dried completely. A further detailed description of *in vitro* phosphosite identification method has been published on protocols.io (dx.doi.org/10.17504/protocols.io.261gen89dg47/v1).

### LC-MS/MS and Data analysis

Vacuum dried peptides of each sample was resolubilized in LC-buffer (3% (v/v) acetonitrile in 0.1% (v/v) formic acid) and incubated on a Thermo mixer at 1800 rpm at room temperature for 30 minutes. Further, 200ng of peptide digest was loaded onto Evotips and samples were prepared as explained in [49]. The Evotips were further placed on an autosampler tray of EvoSep LC system for LC-MS/MS analysis. 30 sample per day (44 min run time) method was used and the peptides were resolved on a 15cm analytical column (ReproSil-Pur C18, 1.9 μm beads by Dr Maisch. #EV1113) and directly electrosprayed using Easy nano LC source into the Orbitrap Exploris 240 (Thermo Fisher Scientific) mass spectrometer. The data was acquired in a Data dependent mode (DDA) targeting top-10 dependent scans. The full MS and MS2 scans were acquired at 120,000 and 15,000 resolution (m/z 200) respectively and measured using Orbitrap mass analyzer. The precursor ions were fragmented using 28% normalized Higher-energy collisional dissociation (HCD). The AGC target for MS1 and MS2 are set at 3E6 and 1E5 ions respectively and maximum ion injection times were set for MS1 and MS2 are 25 and 100 ms. The dynamic exclusion option was enabled and was set for 5 sec durations.

### Database searches

The MS raw data was processed using MaxQuant software suite version 2.0.3.0 [50]. Human LRRK1 sequence from Uniprot was used as a FASTA sequence to search the data. Trypsin+Lys-C, AspN and Chymotrypsin were set as proteases with a maximum of two missed cleavages were allowed. Oxidation of Met, deamidation of Asn/Gln and phosphorylation of Ser/Thr were set as a variable modification and Carbamidomethylation of Cys was set as a fixed modification. The default instrument parameters for MS1 and MS2 tolerance were used, and the data was filtered for 1% PSM, peptide and Protein level FDR. The pSTY sites table was further processed using inhouse python script (https://zenodo.org/record/6627264) to generate the heatmap representation of identified high-confident LRRK1 phosphosites.

### LCMS analysis of LRRK1 digests using ZenoTOF 7600

LRRK1 tryptic digests were analyzed using a Waters M Class UPLC system coupled to ZenoTOF 7600 mass spectrometer (Sciex). Digests were chromatographed on a Kinetix XB C18 150 x 0.3mm column (Phenomenex) at 6 ml/min (A=0.1% Formic acid in water, B=0.1% formic acid in acetonitrile) with a 21min gradient (3-30% B) followed by 3 min (30-80% B). The column was connected directly to the ZenoTOF 7600 OptiflowTM source fitted with a low microflow probe and heated at 30oC with the integral column oven.

LCMS data was acquired using optimal source conditions and the data dependent acquisition was performed on the Top 30 precursors (m/z 400-2000) with charge state 2-4+, with a minimum intensity of 300 cps and excluded for 6s after 1 occurrence. TOFMSMS spectra (m/z 100-2000) were acquired with Electron Activated Dissociation (EAD) using a filament current of 3000nA, 0eV kinetic energy, a reaction time of 10ms and total MSMS accumulation time of 15ms. Data files (wiff) were converted to mgf files using the AB Sciex MS Data converter (Sciex) and searched using Mascot 2.6 against Swissprot (2019_11.fasta) database. Enzyme cleavage allowed for 1 missed cleavage, Carbamidomethylation of cysteine was a fixed modification, phosphorylation of serine, threonine, tyrosine, and oxidation of methionine were variable modifications. The instrument type chosen to match the EAD spectra was EThcD and the mass accuracy for precursors (20ppm) and for MSMS spectra (0.05Da).

## Funding

D.R.A and S.K. laboratories are funded by the joint efforts of The Michael J. Fox Foundation for Parkinson’s Research (MJFF) and Aligning Science Across Parkinson’s (ASAP) initiative. MJFF administers the grant (ASAP-000463-D.R.A. and ASAP-000159-S.K.) on behalf of ASAP and itself. The D.R.A lab is also supported by the UK Medical Research Council [grant number MC_UU_00018/1] and the pharmaceutical companies supporting the Division of Signal Transduction Therapy Unit (Boehringer Ingelheim, GlaxoSmithKline, Merck KGaA.).

## Acknowledgements

We thank Nicole Polinski at the Michael J Fox Foundation for Parkinson’s research for coordinating the generation of the rabbit monoclonal pRab7A Ser72 antibody that made this study possible, and Suzanne Pfeffer (Stanford), Samara Reck-Peterson, Andres Leschziner (UCSD) and Ji Sun (St Jude) for helpful discussions. We also thank the excellent technical support of the MRC protein phosphorylation and ubiquitylation unit (PPU) DNA sequencing service (coordinated by Gary Hunter), the MRC-PPU tissue culture team (coordinated by Edwin Allen), MRC-PPU Reagents and Services cloning team (coordinated by Dr Rachel Toth), antibody and protein purification teams (coordinated by Dr James Hastie), and mass spectrometry team (coordinated by Dr Renata Filipe Soares). For the purpose of open access, the authors have applied a CC BY public copyright license to all Author Accepted Manuscripts arising from this submission.

## Author contributions

A.U.M. and A.K. planned and performed most experimental work, analyzed data, prepared figures, and helped write manuscript. R.S.N. R.G. and N.M. performed the mass spectrometry analysis related to Fig 6B and SFig 3. D.C., S.K., and S.M. generated wild type and mutant recombinant LRRK1, T.K.P. performed data analysis for SFig 3., M.L. performed most of the cloning and D.R.A contributed to conceptualization of project, data analysis, writing manuscript and supervised the work.

## Data availability

All the primary data that is presented in this study has been deposited in Zenodo and repository and can be accessed using the Digital Object Identifier 10.5281/zenodo.6974577. The mass spectrometry proteomics data have been deposited to the ProteomeXchange Consortium via the PRIDE [51] partner repository with the dataset identifier PXD034420. All reagents (and associated datasheets) generated at the MRC Protein Phosphorylation and Ubiquitylation Unit at the University of Dundee can be requested using the indicated DU number identifier through our reagent’s website https://mrcppureagents.dundee.ac.uk/.

---

**SFigure 1:**
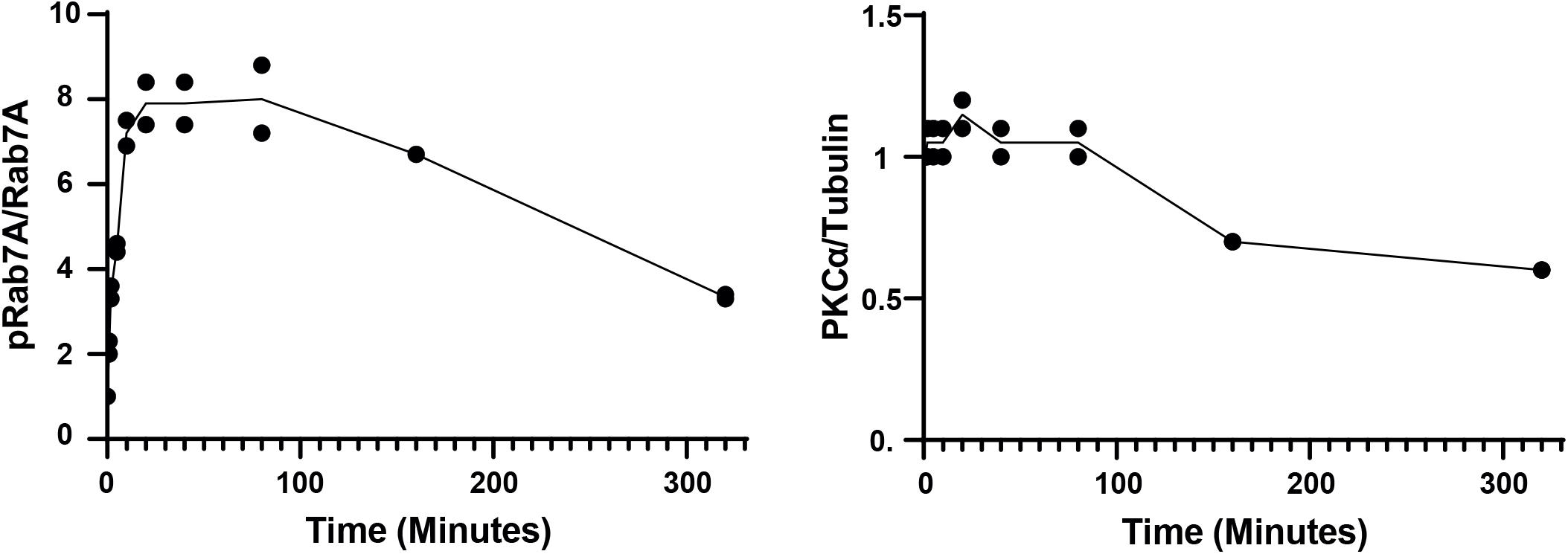
Quantification of time course of PMA treatment in HEK293 Flp-In T-Rex cells. Quantified immunoblotting data from Figure 1C are presented as ratios of pRab7ASer72/total Rab7A and total PKCα/tubulin (mean ± SEM).

**SFigure 2:**
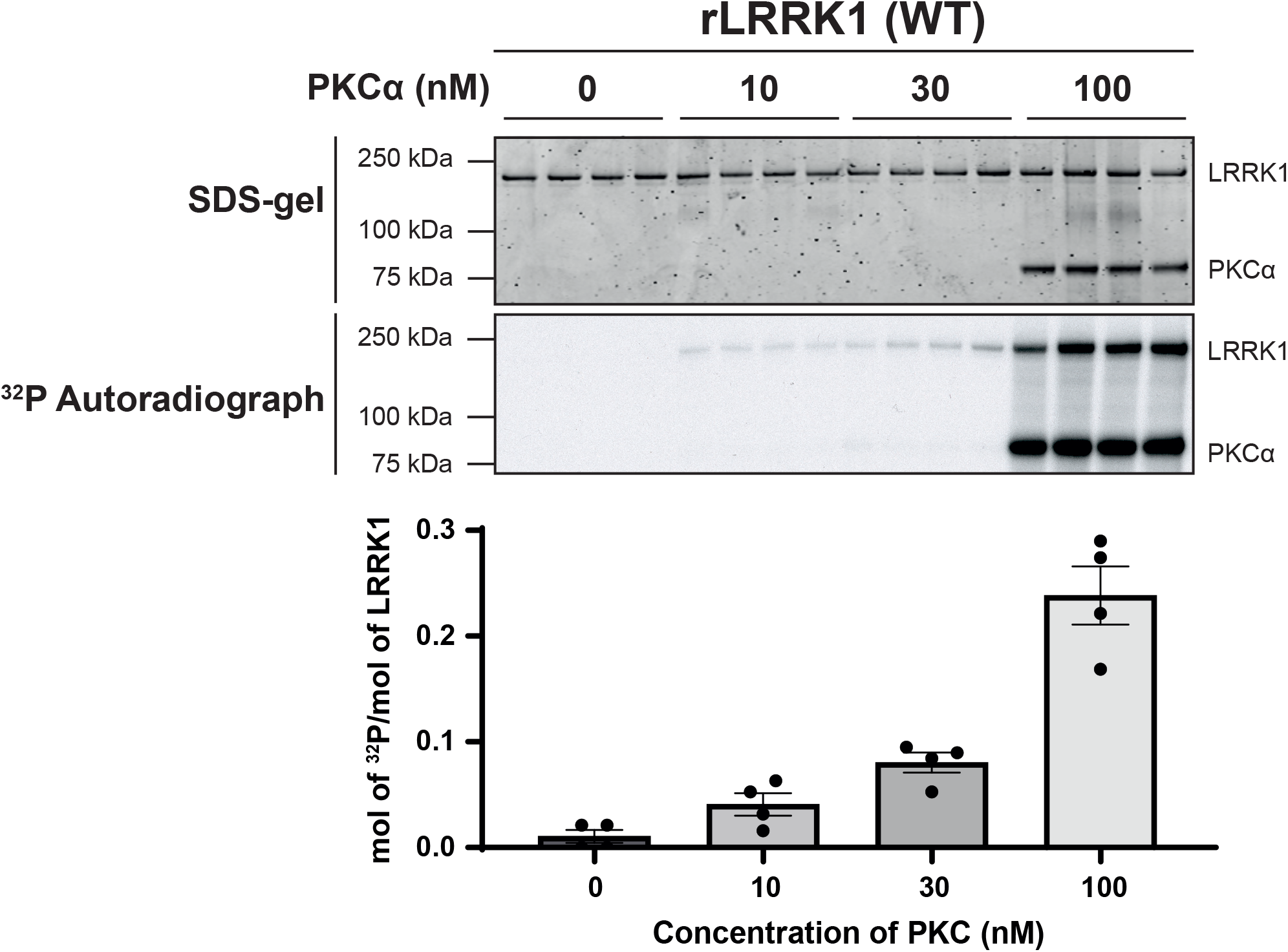
PKCα stoichiometrically phosphorylates LRRK1. The indicated concentrations of PKCα (0-100 nM) was incubated with recombinant wild type (WT) insect cell expressed recombinant (r) LRRK1[20-2015] (50 nM) in the presence of Mg[γ-^32^P]ATP (500 cpm/pmol). Each sample is analysed in quadruplicate. Reactions were terminated after 30 min with SDS-sample buffer. 80% of each reaction was resolved by SDS-polyacrylamide electrophoresis, stained by Coomassie blue (upper panel), and subjected to autoradiography (middle panel). Dried bands were then excised from the gel and counts per minute were evaluated using a scintillation counter to determine stoichiometry of LRRK1 phosphorylation. Data is presented mean ± SEM.

**SFigure 3.**
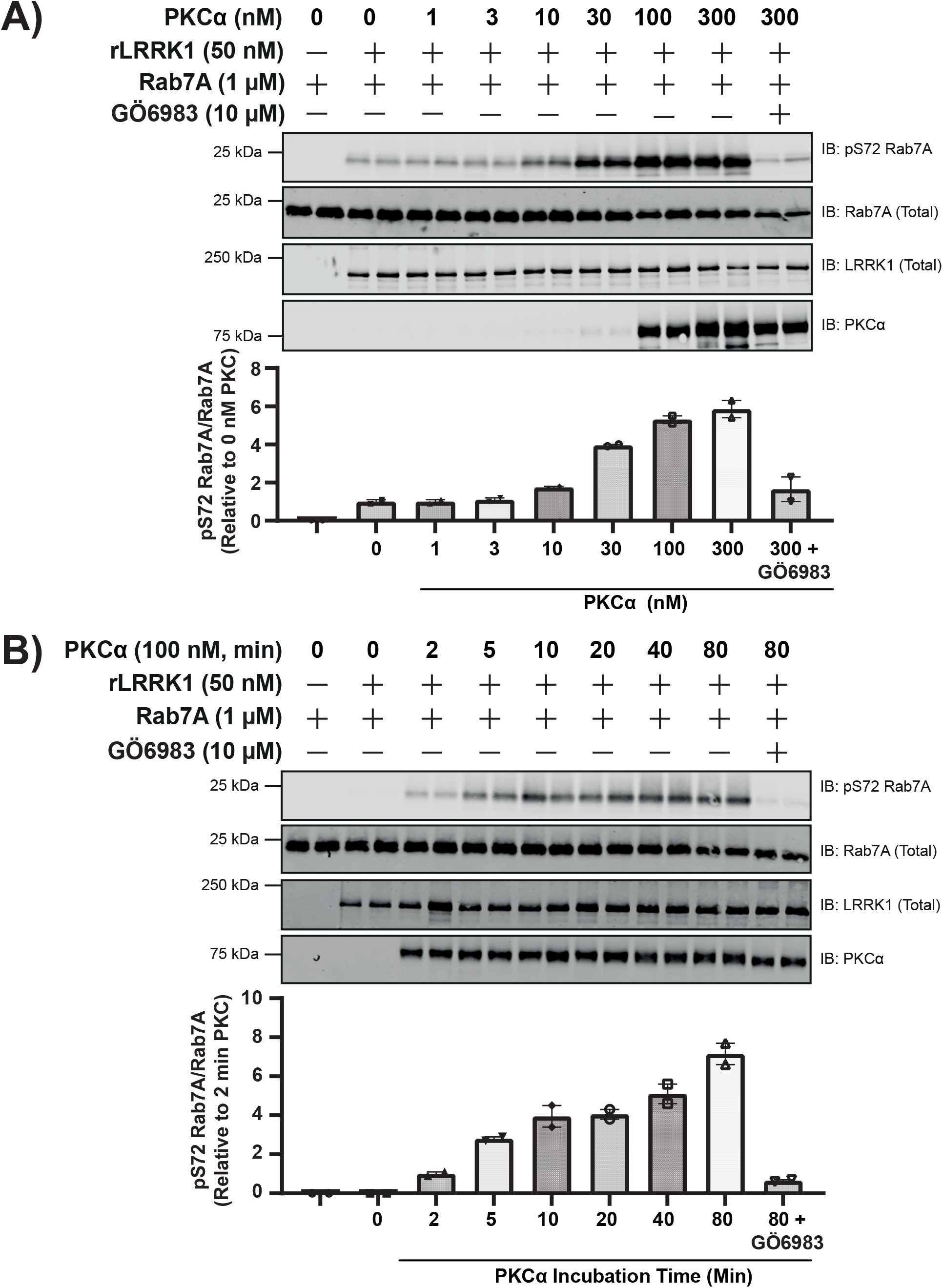
Time-course and dose-dependence of activation of recombinant insect LRRK1 by PKCα. **(A)** Step-1 of the LRRK1 kinase activation assay was setup with insect cell produced recombinant wild type (WT) LRRK1[20-2015], the indicated concentrations of PKCα, ± PKC inhibitor GÖ6983 (10 μM) in the presence of non-radioactive MgATP for 30 min. PKC phosphorylated LRRK1 was then diluted 1.5-fold into a kinase assay containing recombinant Rab7A (1 mM) in the presence of non-radioactive MgATP (Step 2). Reactions were terminated after 30 min with SDS-sample buffer and subjected to a multiplexed immunoblot analysis using the LI-COR Odyssey CLx Western Blot imaging system with the indicated antibodies. Combined immunoblotting data from 2 independent biological replicates (each performed in duplicate) are shown. Lower panel quantified immunoblotting data are presented as ratios of pRab7ASer72/total Rab7A (mean ± SEM) relative to levels observed with no PKCα added (give a value of 1.0). (**B**) As in (**A**) except in step 1 of the kinase activity assay 100 nM PKCα was incubated with immunoprecipitated GFP-LRRK1 for the times indicated.

**SFigure 4:**
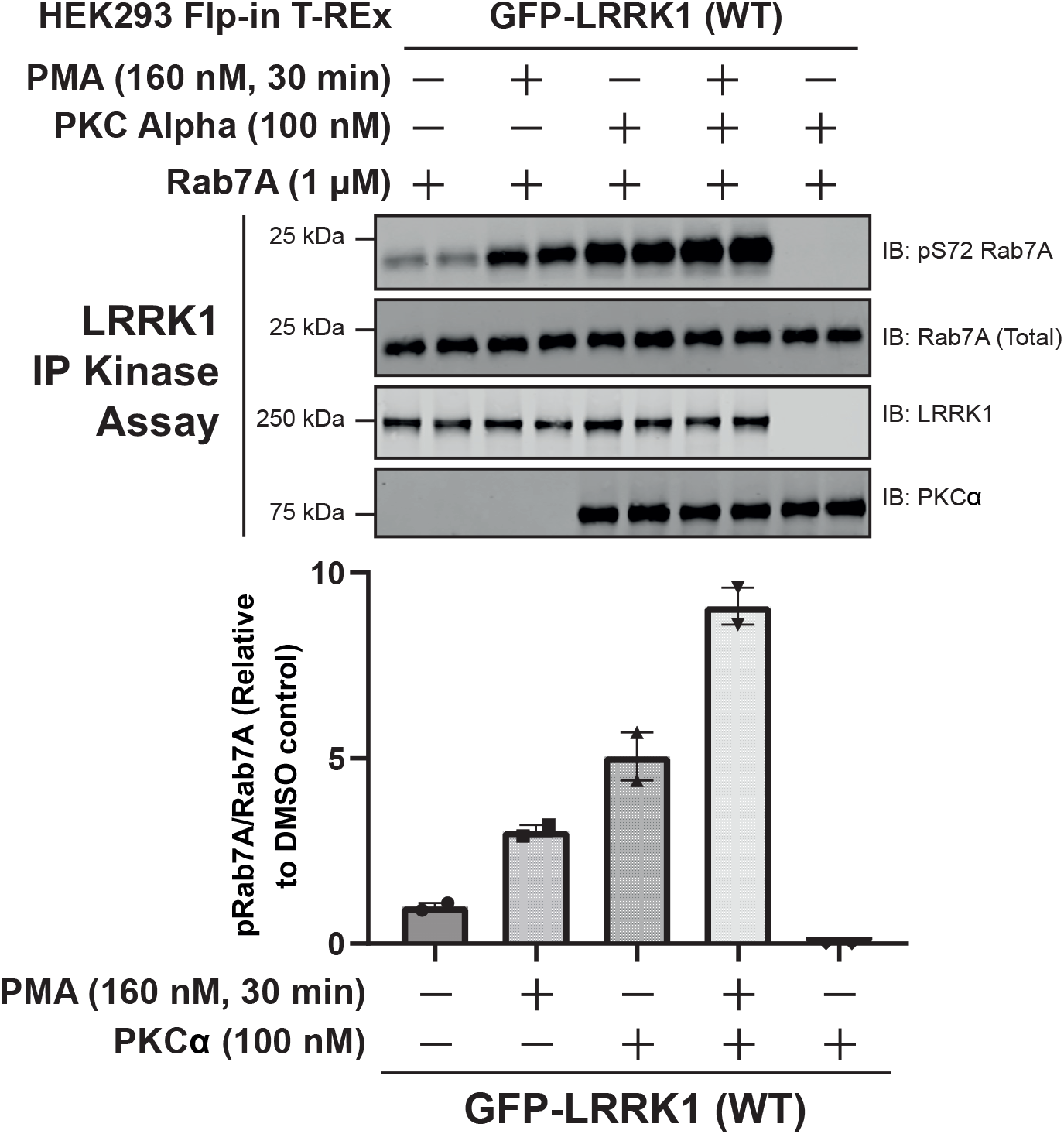
Recombinant PKC Alpha further activates PMA-stimulated LRRK1. HEK293 Flp-In T-REx cells stably expressing wild type (WT) GFP-LRRK1 were treated ± 1 μg/ml doxycycline for 24 h to induce GFP-LRRK1 expression. Cells were serum-starved for 16 h and stimulated ± 160 nM phorbol 12-myristate 13-acetate (PMA) for 30 min and lysed. GFP-LRRK1 immunoprecipitated and aliquoted as indicated. Each aliquot contains GFP-LRRK1 immunoprecipitated from 1 mg of HEK293 cell lysate. Step-1 of the LRRK1 kinase activation assay was setup by incubating GFP-LRRK1 aliquots with PKCα (100 nM) in the presence of non-radioactive MgATP for 30 min. PKC phosphorylated LRRK1 was then diluted 1.5-fold into a kinase assay containing recombinant Rab7A (1 mM) in the presence of non-radioactive MgATP (Step 2 of the kinase assay). Reactions were terminated after 30 min with SDS-sample buffer analysed by multiplexed immunoblot analysis using the LI-COR Odyssey CLx Western Blot imaging system with the indicated antibodies (upper panel). Combined immunoblotting data from 2 independent biological replicates (each performed in duplicate) are shown. Lower panel quantified immunoblotting data are presented as ratios of pRab7ASer72/total Rab7A (mean ± SEM) relative to levels observed with no PKCα added (given a value of 1.0).

**SFigure 5.**
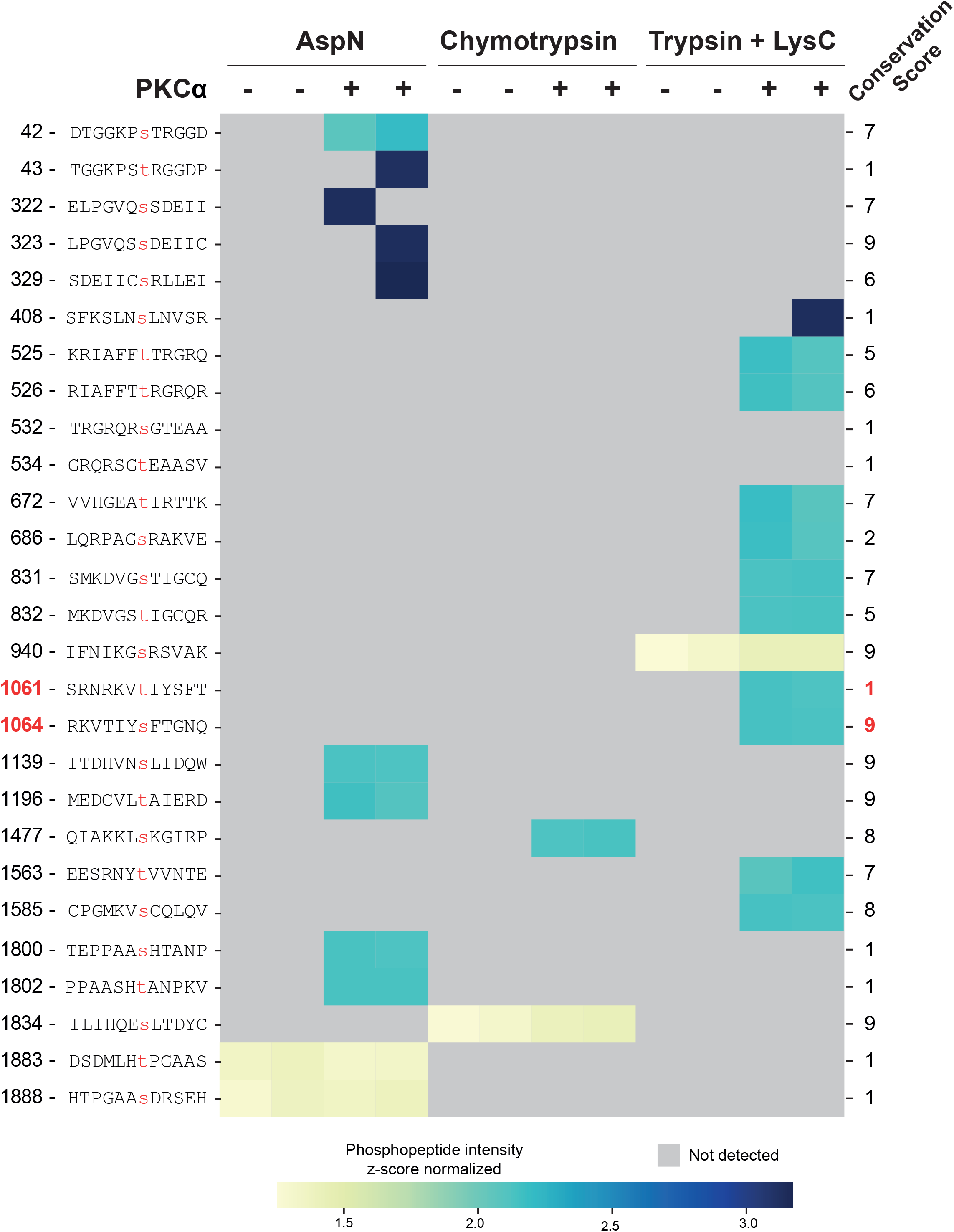
Heatmap of phosphorylation sites identified via HCD analysis. Kinase inactive LRRK1[D1409A, 20-2015] (200 nM) was incubated ± PKCα (400 nM) the presence of MgATP. Reactions were terminated after 30 min with SDS-sample buffer and reactions resolved by SDS-polyacrylamide electrophoresis and gel stained by Coomassie blue. The gel bands containing LRRK1 were digested with mixture of AspN, Chymotrypsin and Trypsin+Lys-C. The resultant peptides were analyzed by HCD fragmentation acquired on Orbitrap Exploris 240 MS platform. The high confidence phosphosites that are identified and analyzed using MaxQuant are depicted as a heatmap representing the relative abundance of identified phosphosites versus protease utilized ± PKCα. The grey color denotes no phosphopeptides detected. Side panel show the evolutionary conservation score from 1 (least conserved) to 9 (most conserved) of each of the identified phosphorylation sites determined calculated the Consurf server ((https://consurf.tau.ac.il/) [52]. Sequences of LRRK1 homologs were obtained from OrthoDB (https://www.orthodb.org/) [53] and alignment of sequences was performed using the MAFFT server (https://www.ebi.ac.uk/Tools/msa/mafft/) [54]. Phosphosites Thr1061 and Ser1064 and their respective conservation scores are highlighted in red.

**SFigure 6.**
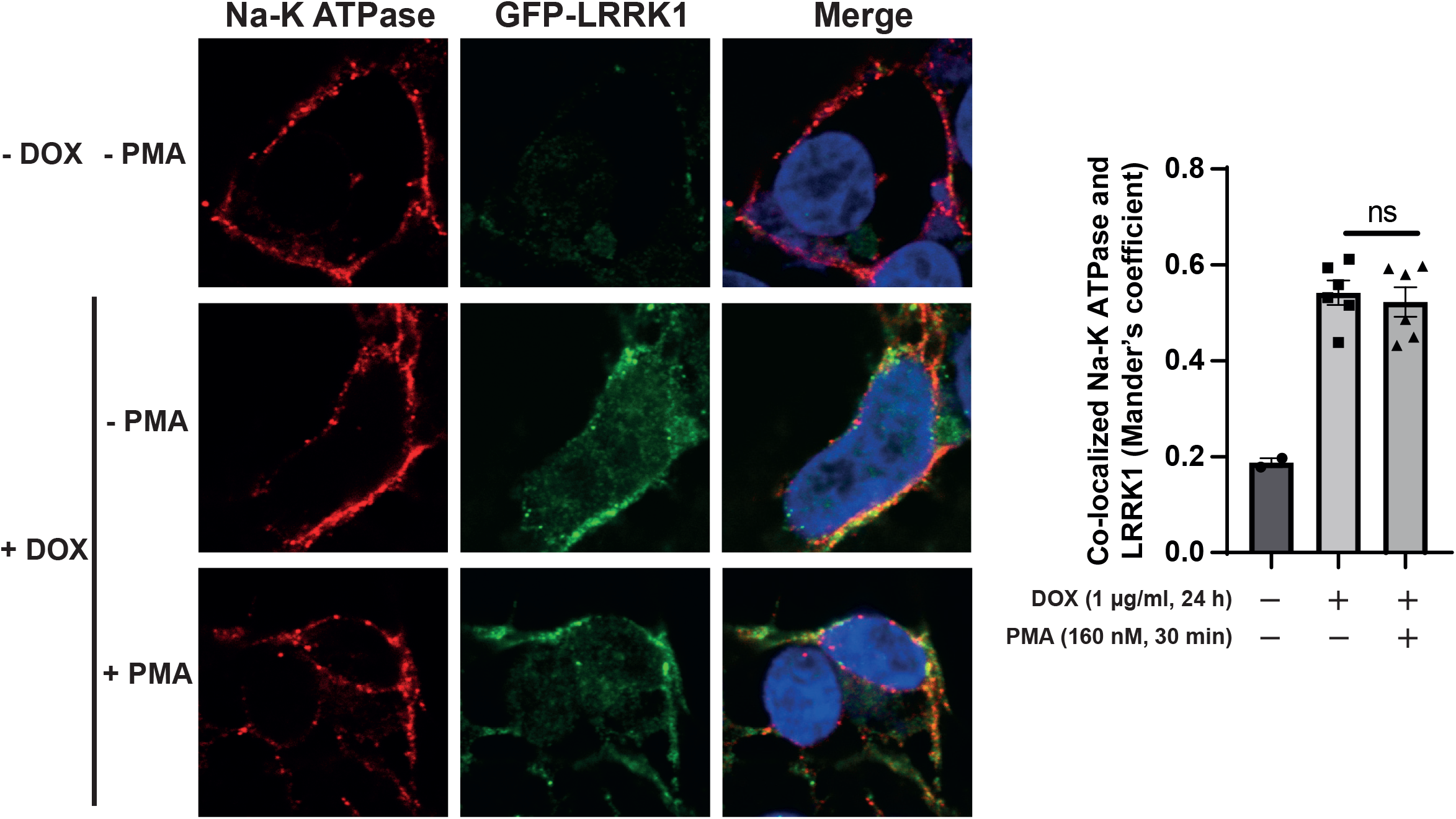
Membrane-fraction of LRRK1 colocalizes with plasma membrane marker Na-K ATPase. **Left panel)** HEK293 Flp-In T-REx cells stably expressing wild type (WT) GFP-LRRK1 were treated with 1 μg/ml doxycycline for 24 h to induce GFP-LRRK1 expression. Cells were serum-starved for 16 h and stimulated ± 160 nM phorbol 12-myristate 13-acetate (PMA) for 30 min. Cells were permeabilized by liquid nitrogen freeze–thaw to deplete cytosol [48] and then fixed and stained with mouse anti-Na-K ATPase, chicken anti-GFP and DAPI. **(Right panel)** Colocalization of GFP-LRRK1 and Na-K ATPase was determined from a Mander’s coefficient (presented as mean ± SEM) after automatic thresholding. *P* = 0.6425 (ns) by Student’s unpaired, two-tailed t test.

**SFigure 7.**
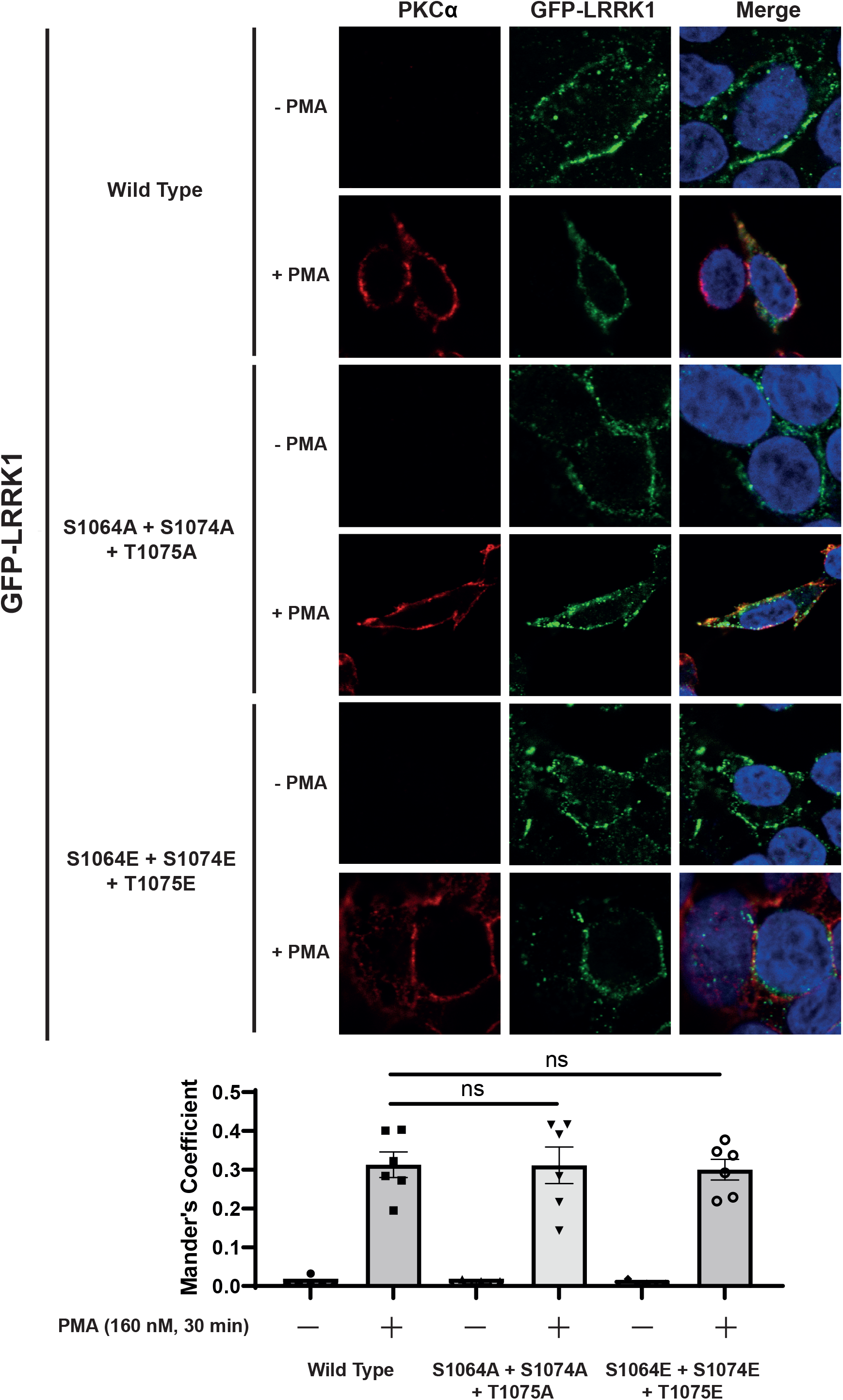
Triple Ser1064, Ser1074 and Thr1075 alanine or glutamate mutations do not affect LRRK1 localization. (Left panel) HEK293 cells were grown on cover slips and transiently transfected with plasmids encoding GFP-LRRK1 wild type (WT) or the indicated mutants of GFP-LRRK1. Cells were serum-starved for 16 h and stimulated ± 160 nM phorbol 12-myristate 13-acetate (PMA) for 30 min. Cells were permeabilized by liquid nitrogen freeze–thaw to deplete cytosol [46] and then fixed and stained with mouse anti-PKCα, chicken anti-GFP and DAPI. (Right panel) Co-localization of GFP-LRRK1 and PKCα, was determined from a Mander’s coefficient (presented as mean ± SEM) after automatic thresholding. P > 0.05 (ns) by Student’s unpaired, two-tailed t test.

## Notes

### Competing Interest Statement

The authors have declared no competing interest.

### Summary of Updates

This version of the manuscript has been revised to address Reviewers comments and resubumitted for publication in the Biochemical Journal

